# Doubled haploid based parental lines are most suitable in predicting heterosis using microsatellites and in development of highly heterotic F_1_ hybrids in *Brassica oleracea*

**DOI:** 10.1101/511055

**Authors:** Saurabh Singh, S.S. Dey, Reeta Bhatia, Raj Kumar, Kanika Sharma, T.K. Behera

## Abstract

In *Brassica oleracea*, heterosis is one of the most efficient tools giving impetus to hybrid vegetable industry. In this context, we presented the first report on identifying superior heterotic crosses for yield and commercial traits in cauliflower involving cytoplasmic male sterile (CMS) and doubled haploid (DH) lines as parents. We studied the suitability of SSR and EST-SSRs based genetic distance (GD) and morphological markers based phenotypic distance (PD) in prediction of heterosis when DH based genotypes are used as parents in developing F_1_ hybrids. Overall 120 F_1_ hybrids derived from twenty *Ogura* cybrid CMS lines and six DH based testers were evaluated for 16 phenotypic traits along with their 26 parental lines and 4 commercial standard checks, in 10 × 15 alpha lattice design. The genomic SSR and EST-SSRs based genetic structure analysis grouped 26 parental lines into 4 distinct clusters. The CMS lines Ogu118-6A, Ogu33A, Ogu34-1A were good general combiner for developing short duration hybrids. The SCA effects were significantly associated with heterosis suggesting non-additive gene effects for heterotic response of hybrids. Less than unity value of σ^2^A/D coupled with σ^2^_gca_/σ^2^_sca_indicated the predominance of non-additive gene action in the expression of studied traits. The genetic distance estimates among 26 parents ranged from 0.44 to 0.98 and were significantly associated with heterosis for important commercial traits, suggesting the utility of microsatellite based genetic distance in prediction of heterosis in *B. oleracea*.

## Introduction

In the plant kingdom, the family *Brassicaceae* holds a great agronomic, scientific and economically important position and comprises more than 372 genera and 4060 species [2]. Of the diverse species, *Brassica oleracea* (CC, 2n = 18) constitutes well defined group of economically and nutritionally important morphotypes, referred to as cole vegetables (kale, kohlrabi, cabbage, cauliflower, broccoli, brussels sprout) [52]. These *Brassica* vegetables are also termed as ‘super-food’ as they are vital source of secondary metabolites, antioxidants, vitamins and minerals [71, 16, 64, 69]. Among the cultivated *B. oleracea* varieties, cauliflower (*B. oleracea* var. *botrytis* L.) is an important vegetable crop grown worldwide. Great efforts have been made to improve the productivity and quality of this crop, ascribed to its economic value and as an essential component of healthy diet [91]. The replacement of open pollinated varieties with F_1_ hybrids have become much pronounced in cole vegetables including cauliflower due to high uniformity, better quality, tolerance to various biotic stresses and vagaries of adverse climatic conditions [17, 69]. It is well established that *Brassica* vegetables exhibit a wide range of heterosis and high heterosis have been reported in cauliflower also for both yield and quality traits [13, 15, 69]. Nature has bestowed *Brassica* vegetables with genetic mechanisms of sporophytic self-incompatibility (SI) and cytoplasmic male-sterility (CMS), which have efficiently triggered the hybrid breeding programme in these crops [13, 15, 17, 66, 69, 77]. However, in the current scenario of increasing temperature as a result of global warming there is frequent breakdown of self-incompatibility, as S-alleles are more prone to high temperature. Thus, SI lines are not always stable and results in sibbed seed in hybrid breeding [66]. In addition, the maintenance of S-allele lines is time consuming and costly endeavor and in case of snowball cauliflower, SI system is very poor or absent [66, 68]. Under these circumstances, the genetic system of CMS provides a better alternative for heterosis breeding in cole crops [17, 69]. Heterosis or hybrid vigor, plausibly results from accumulation of parental genetic and epigenetic information, is manifested as superior performance of hybrid offspring relative to the average of their genetically diverse parents [3, 26, 45]. Despite its tremendous economic value in hybrid breeding, the molecular basis behind this biological phenomenon is still obscure [3, 49, 26, 31, 48]. So far, different hypothesis and genetic mechanisms have been put forward to elucidate this complex phenomenon such as dominance model, over-dominance model and epistasis model [3, 49, 26, 31, 48]. Recent progress in QTL analysis, transcriptomics, proteomics, metabolomics have helped in elucidation of heterosis at molecular level to some extent by explaining the role of epigenetic regulations such as DNA methylation, small RNAs (sRNAs) and histone modifications in hybrid vigour in different crop plants [26, 31, 45, 48]. Preselecting inbred parents and recognizing most promising heterotic combinations is crucial for accelerating heterosis breeding in crop plants. The measure of both general combining ability (GCA), which provides information on breeding value of parents as well as about the additive genetic control, and specific combining ability (SCA) is necessary for selection of parental lines and heterotic groups [38]. The estimation of SCA is associated with non-additive effects (dominance effects, additive×dominant and dominant×dominant interactions). Among different biometrical approaches, line × tester analysis appears to be ideal for estimating GCA effects of lines and testers, SCA effects of cross combinations, and providing information about nature of gene actions [17, 20, 41]. The extent of heterosis has been reported to be vary with mode of reproduction, genetic distance of parents, traits under investigation, developmental stage of plant and prevailing environment [27, 34, 38, 40, 44, 73]. The pair-wise parental genetic distance (GD) has been suggested as a good indicator of *per se* hybrid performance and recognition of heterotic groups [27, 34, 38, 40, 44, 73]. Different approaches are available to determine genetic distance depending upon morphological traits, horticultural data, biochemical characteristics and DNA markers based genotypic data [33, 55]. Molecular markers have been well established as a powerful tool for analyzing genetic diversity and estimation of genetic distances among different genotypes or advance breeding populations. SSR (simple sequence repeat) markers have been markers of choice owing to their co-dominant inheritance, whole-genome coverage, abundance and high reproducibility [40, 76]. However, so far, contradictory results have been reported with respect to relationship between GD and heterosis across different crops (3, 20, 27, 33, 34, 36, 40, 44, 73, 78]. These results suggest that the heterosis for yield and yield related traits is highly complex phenomenon. As suggested by Cress [12] that for significant heterosis the extent of parental GD is essential but is not enough to assure it and in addition, the better forecasting of heterosis is possible only when GD is lesser than a definite threshold level [4]. Furthermore, the association of GD and heterosis also depends upon the germplasm, population under investigation, methods of calculating GD [73-74]. The parents with small GD can also display high level of heterosis like closely related ecotypes in *Arabidopsis* resulted hybrids with significant improvement in plant biomass and seed yield [26, 32]. Contrasting results are also available about association of genetic distance based on morphological traits (hereafter referred as PD: phenotypic distance) and mid-parent heterosis (MPH) and SCA in various crop plants [33, 78, 79, 86]. Teklewold and Becker [79] reported significantly positive association of PD with MPH, GCA and hybrid performance for seed yield in Ethiopian mustard (*Brassica carinata*), while Hale et al. [34] found no correlation of PD with heterosis in broccoli.

The development of homozygous inbred lines is tedious and time consuming process in *B. oleracea* crops due to their allogamous nature, on account of genetic mechanisms of protogyny and self-incompatibility [9] leading to high inbreeding depression. In the heterosis breeding programmes, the inbred development is prerequisite for successful hybrid development programmes and for different genetic studies. The availability of doubled haploid (DH) technology eliminates the long time requirement for inbred generation through traditional 5-7 generations of selfing. The DH technology in *B. oleracea* crops through isolated microspore culture (IMC) has accelerated the breeding programmes through generation of homozygous DH lines in two-successive generations [9, 25]. The DH induction has significantly enhanced the genetic and genomic research in *Brassica* vegetables. The DH based breeding populations have been instrumental in discovery and mapping of QTLs of economically important agronomic and quality traits in *Brassica* vegetables [35, 51, 72, 89]. DH based populations has also facilitated the construction of high-density genetic linkage map [90], mapping QTLs/genes for disease resistance [50, 70], identification of QTLs related to timing of curd induction, subtropical adaptation in *Brassica oleracea* crops [35, 46, 62]. Thus, realizing the utility of DH technology in accelerating the genetic improvement of *B. oleracea* crops, completely homozygous DH lines have been developed by our group previously through IMC in cauliflower [7-9]. Concurrently, advance generation *Ogura* CMS lines in cauliflower for heterosis breeding have also been developed by protoplast fusion followed by recurrent backcrossing in the nuclear background of elite genotypes [6, 13].

To the best of authors’ knowledge, rare or inadequate information is available regarding combining ability, gene action and heterosis breeding in cauliflower particularly using CMS and DH system for yield and agro-morphological traits. Recently, we have reported heterotic responses by combining ability analysis utilizing this combination of CMS and DH system for antioxidant capacity and quality traits in cauliflower [69]. Then, to our knowledge we have not found any study in this particular crop (*B. oleracea* var. *botrytis* L.) about association of molecular GD and morphological PD with MPH, beter-parent heterosis (BPH) and SCA for yield and commercial traits. Thus, if genetic distance is significantly correlated with heterosis for commercial traits in cauliflower, the parental selection could be done based on genetic distance instead of field trials. Although, contrasting results have been obtained in different crops in this context, as heterosis is complex biological phenomenon with numerous genes numerous genetic mechanisms (26, 31, 45, 48]. Hence, in the present investigation, the main objectives were to (i) identify heterotic groups of CMS and DH lines for hybrid breeding on the basis of GCA, SCA effects, nature of gene action and heritability in cauliflower (ii) to find out is there any correlation of SSRs, EST-SSRs (expressed sequence tag based-SSRs) based GD and morphological traits based PD with heterosis and SCA (iii) to investigate the association of SCA with MPH, BPH and also to study the SSR and EST-SSRs based population structure of parental lines and testers used in the study. Present investigation is the first report of heterosis and combining ability based on CMS and DH technique in cauliflower to examine the prospects of developing F_1_ hybrids with respect to yield and commercial traits and to assess the role of genetic distances in prediction of heterosis in cauliflower.

## Materials and methods

### Plant materials, experimental site, mating and experimental design

The field experiment was carried out at Baragram Experimental Farm of ICAR-Indian Agricultural Research Institute (IARI), Regional Station, Katrain, Kullu Valley, Himachal Pradesh, India. The experimental farm is located at 32.12N latitude and 77.13E longitudes with an altitude of 1,560 m above mean sea level. The basic genetic plant material for the present investigation comprised of 20 genetically diverse *Ogura* cybrid cytoplasm based elite CMS lines previously developed after more than nine generations of backcrossing having desirable agronomic and floral traits (Table 1). These CMS lines were used as female parent in the breeding programme. The completely homozygous 6 DH inbred lines of snowball cauliflower with abundant pollen production, developed through IMC, were used as testers (Table 1).

**Table 1.** Parental lines (cytoplasmic male-sterile) and testers (doubled haploid) used in the study

| Code | Line | Curd Color | Curd compactness | Curd covering by inner leaves | Riceyness | Anthocyanin pigmentation |
| --- | --- | --- | --- | --- | --- | --- |
| L1 | Ogu122-5A | White | Compact | PC | Absent | Absent |
| L2 | Ogu115-33A | White | Compact | PC | Absent | Absent |
| L3 | Ogu118-6A | White | Compact | PC | Absent | Absent |
| L4 | Ogu307-33A | Creamy White | Compact | NC | Absent | Absent |
| L5 | Ogu309-2A | Creamy White | Compact | PC | Absent | Absent |
| L6 | Ogu33A | White | Compact | FC | Absent | Absent |
| L7 | OguKt-2-6A | White | Compact | PC | Absent | Absent |
| L8 | Ogu1A | White | Compact | PC | Absent | Absent |
| L9 | Ogu13-85-6A | White | Compact | NC | Absent | Absent |
| L10 | Ogu1-6A | White | Compact | PC | Absent | Absent |
| L11 | Ogu2A | White | Compact | PC | Absent | Absent |
| L12 | OguKt-9-2A | White | Compact | PC | Absent | Absent |
| L13 | Ogu22-1A | Creamy White | Compact | PC | Absent | Absent |
| L14 | Ogu122-1A | White | Compact | PC | Absent | Absent |
| L15 | Ogu126-1A | White | Compact | PC | Absent | Absent |
| L16 | Ogu12A | White | Compact | PC | Absent | Absent |
| L17 | Ogu119-1A | Creamy White | Compact | PC | Absent | Absent |
| L18 | Ogu34-1A | White | Compact | PC | Absent | Absent |
| L19 | Ogu125-8A | White | Compact | FC | Absent | Absent |
| L20 | Ogu33-1A | Creamy White | Compact | PC | Absent | Absent |
| T1 | *DH-18-8-1 | White | Compact | PC | Absent | Absent |
| T2 | *DH-18-8-3 | White | Compact | PC | Absent | Absent |
| T3 | *DH-53-1 | White | Compact | FC | Absent | Absent |
| T4 | *DH-53-6 | White | Compact | FC | Absent | Absent |
| T5 | *DH-53-9 | White | Compact | PC | Absent | Absent |
| T6 | *DH-53-10 | White | Compact | PC | Absent | Absent |

\*All the DH lines used as testers were developed through isolated microspore culture (IMC) and assessment of their ploidy level through flow cytometry analysis; L: lines, T: testers, PC: partly covered, FC: fully covered, NC: not covered

The CMS and DH lines were selected among the 60 CMS and 24 DH lines developed previously, based on molecular, morphological characterization (data not shown in this publication) and flowering synchronization of lines and testers was also the main consideration in selection of parents. All the recommended package of practices, suggested for raising cauliflower crop at IARI- regional station Baragram farm, were followed to grow a healthy crop for displaying better agronomic and phenotypic expression [67]. The size of plot was kept 3.0 x 3.0 m^2^ with inter-and intra-row spacing of 45 cm. Then following the line x tester mating design [41], 20 CMS lines of cauliflower were crossed with 6 DH male fertile testers at flowering to generate 120 test cross progenies. To avoid any natural pollination, CMS lines were grown under muslin cloth cage. To pollinate fully opened flowers of CMS lines, fresh pollen from DH testers grown under net house was collected. Each CMS line was pollinated with all six DH testers for the hybrid seed production. Then, healthy seedlings of all the 120 F_1_ hybrids and their 26 parental lines (20 CMS + 6 DH) along with 4 commercial CMS based hybrids (HVCF-18, HVCF-29, HVCF-16 from Acsen HyVeg and Pahuja from Pahuja Seeds) as standard checks, were transplanted at the Baragram Experimental Farm of IARI to evaluate them for morphological, horticultural and yield related traits. All the 120 testcross progenies along with their parents and commercial checks were evaluated in 10×15 alpha lattice experimental design with three replications. For data recording of agronomic traits, five randomly selected well established plants were tag-labelled in each plot/block/replication.

### Morphological and agronomical characterization

The 20 *Ogura* CMS lines, 6 DH testers along with their 120 test cross progenies were evaluated for sixteen agro-morphological traits viz. (i) days to 50% curd initiation: Days to 50% CI) (ii) days to 50% curd maturity: Days to 50% CM (iii) plant height: PH (cm) (PH) (iv) gross plant weight: GPW (g) (v) marketable curd weight: MCW (g) (vi) net curd weight: NCW (g) (vii) leaf length: LL (cm) (viii) leaf width: LW (cm) (ix) number of leaves: NoL (x) curd length: CL (cm) (xi) curd diameter: CD (cm) (xii) core length: CoL (cm) (xiii) curd size index: CSI (cm^2^) (xiv) leaf size index: LSI (cm^2^) (xv) harvest index: HI (%) (xvi) Total marketable yield: TMY (t/ha) [13-15, 17]. Data were recorded from 5 randomly selected plants of each genotype of each plot/block of all the three replications.

### Statistical analysis for agronomic traits

The agronomic data recorded for each parent, 120 F_1_ hybrids and 4 commercial checks in alpha lattice design were subjected to analysis of variance (ANOVA) using GLM procedure of SAS (statistical analysis system) software version 9.4 [65]. The line × tester statistical analysis of GCA, SCA, heterosis, heritability, variance and mean performance for was accomplished as per Kempthorne [41] through SAS version 9.4. The testing of significance of GCA and SCA effects was done at 5%, 1%, and 0.1% probability through F test. Heterosis estimates for different traits were computed as per Xie et al. [85] based on formulae viz, MPH% (Mid parent heterosis) = [(F_1_-MP)/MP] x 100, BPH% (Better parent heterosis) = [(F_1_-BP)/BP] x 100, where MP is mid-parent and BP is better-parent performance and testing of significance was done at probability of p < 0.05, p < 0.01 and p < 0.01 through *F* test. The narrow-sense heritability (h^2^_ns_ = V_A_/V_P_; V_P_ = V_G_ + V_E_) estimates were categorized into three classes viz., high (> 30%), medium (10-30%) and low (< 10%) [61]. The GA was calculated as = H^2^_b_ x phenotypic standard deviation x K, where K value is 2.06, which is standardized selection differential constant at 5% selection intensity [37]. The parental lines and testers were clustered into different groups based on sixteen agronomic traits using R software [63]. They were grouped through principal component analysis (PCA) to estimate the explained variance in first two axes. Pooled data from five randomly selected plants of each genotype per plot per block per replication for all the sixteen morphological and commercial traits were taken for statistical analysis.

### DNA extraction, PCR amplification

All the parental CMS lines and DH testers were grown in pro-trays under glass house conditions in a soilless mixture of cocopeat, perlite and vermiculite in the ratio of 3:1:1. Genomic DNA extraction and purification was done from 100 mg fresh green young expanding leaves of 25-30 days old seedlings using cetyltrimethyl ammonium bromide (CTAB) method with slight modifications [57]. Genomic DNA samples were adjusted to 25-50 ng DNA/µl and also stored at −80 ºC as safeguard for further requirement. For the genotyping purpose, the pair of 350 microsatellite primers comprising genomic-SSRs and EST-SSRs distributed throughout the *Brassica oleracea* genome [47, 82] was used for genetic diversity analysis in parental CMS and DH lines of cauliflower. Among these 145 microsatellite primers were found to be polymorphic and of which 87 SSRs and EST-SSRs displaying clear amplification and polymorphism were used for final molecular analysis of 26 CMS and DH lines. Eppendorf Mastercycler Nexus GSX1 was used for PCR amplification in a reaction volume of 25 μl. The PCR reaction mixture comprised of 1 μl of each forward and reverse primers, 2 μl of genomic DNA template (50 ng), 12.50 μl of 2× PCR Green master mix (GoTaq DNA polymerase; Promega, USA) and 8.50 μl nuclease free water. The PCR cycling programme was set up as follow: an initial denaturation of 94 ºC for 4 min followed by 35 cycles of denaturation at 94 ºC for 30s, annealing of primers at 50 to 60 ºC for 30s depending upon appropriate primer annealing temperatures and extension at 72 ºC for 1min, then final extension of 72 ºC for 7min. Amplified PCR products were separated by 3.0% agarose gel electrophoresis in 1X TBE buffer (pH 8.0) and gel was run at 100 mA voltage for 120 min. Ethidium bromide (EtBr) of 0.5 mg/ml was used for gel staining and gel pictures were captured using one digital gel documentation unit (BioSpectrum^®^ Imaging System^™^, UK). The determination of fragment sizes were done using Promega^TM^ 50 bp DNA step ladder.

### Molecular analysis and genetic structure analysis

Among 350 microsatellite markers, the 87 polymorphic genomic-SSR and EST-SSRs loci depicting genetic diversity (S1 Table) were used for cluster analysis, dendrogram construction based on simple matching (SM) coefficient with the PCA and neighbor joining (NJ) UPGMA method using DARwin software version 6.0.017 [59]. For testing the reliability of NJ dendrogram, a bootstrap value of 1000 replicates was used. For the allelic diversity analysis estimating observed number of alleles (N) per loci, observed heterozygosity (H_o_), expected heterozygosity (H_e_) and polymorphism information content (PIC) were computed through software CERVUS version 3.0 [39]. The estimation of PIC for each locus using CERVUS 3.0 was calculated according to formula; PIC = 1-Ʃ P*i*^2^, where P*i* represents the *i*th allele frequency in a locus for the genotypes *P* under study [53].

The genetic structure analysis of parental population of testcross progenies was studied with Bayesian model-based clustering approach implemented in STRUCTURE version 2.3.4 software [60] to assign individuals to *k* clusters and sub-clusters. For the estimation of proportion of ancestral contribution in each parental line, all simulations were performed by parameter setting as: “admixture model” with “correlated allele frequencies”. The algorithm was implemented with 10,000 length of burn-in period followed by 100000 Markov Chain Monte Carlo (MCMC) repetitions and plausible range of putative k values was kept from k = 1 to k = 10 run independently with 15 iterations for each k. The optimum value of k for determining most likely number of subpopulations was predicted according to simulation method of DeltaK (*ΔK*) [21] with the help of web-based STRUCTURE HARVESTER version v0.6.94 [18].

### Correlation among genetic distances, heterosis, combining ability

The Euclidean distance (ED), hereafter referred as phenotypic distance (PD) was calculated based on sixteen agronomic traits (days to 50% CI, days to 50% CM, PH, GPW, MCW, NCW, LL, LW, NoL, CL, CD, CoL, CSI, LSI, HI, TMY) using R software [63]. The SM dissimilarity coefficient (hereafter referred as genetic distance: GD) was computed based on SSR and EST-SSRs data analysis using DARwin software version 6.0.017. The association among GD, PD, MPH, BPH, SCA was computed by Pearson’s correlation coefficients (r) (pearson product moment correlation coefficient: PPMCC) by using R software pakages version 3.5.1 in Rstudio 1.1.456 [63] and testing of significance at p < 0.05 and p < 0.01. The corrplot displaying correlation among distances, heterosis and combining ability was demonstrated via Rcorrplot package in Rstudio [84].

## Results

### Analysis of variance

The mean square estimates for different vegetative and commercial traits in alpha lattice experimental design revealed significant differences among treatments for all the characters except CD, CoL and CSI at 0.01% probability (Table 2). Likewise, the significant block effects in each replication were found for all the studied traits except Days to 50% CI, CD, CoL and CSI at the probability of 0.01% (Table 2). The coefficient of determination (R^2^) indicated high variability percentage (>70%) for all the traits except except CL, CD, CoL and CSI in the response ascribed to given independent variables (Table 2). The higher R^2^ value also suggests a higher significance of model. The line × tester analysis of variance (ANOVA) for combining ability revealed highly significant differences (P < 0.001) among the treatments and parents for all the vegetative and commercial traits (Table 3) except for CL for which significant differences among parents were found at P < 0.05. The mean squares of lines were also found significant for all the traits at 0.1% except for CL for which significant level of probability was P < 0.05; while the mean squares of testers were non-significant for CL except all other traits (Table 3). The significant differences were also found with respect to lines versus testers for all the traits except LW, CL, CD, CoL, CSI and HI, while the mean squares of parents versus crosses were significant for all the traits except NoL (Table 3). The variance analysis for combining ability also revealed highly significant differences among 120 testcross progenies for all the 16 traits at 0.1% probability, while no significant differences were found among three replications for all the traits except LW, suggesting true presence of inherent variability among all the crosses (Table 3). The line × tester interaction effects were also significant for all the 16 agronomic traits.

**Table 2.**
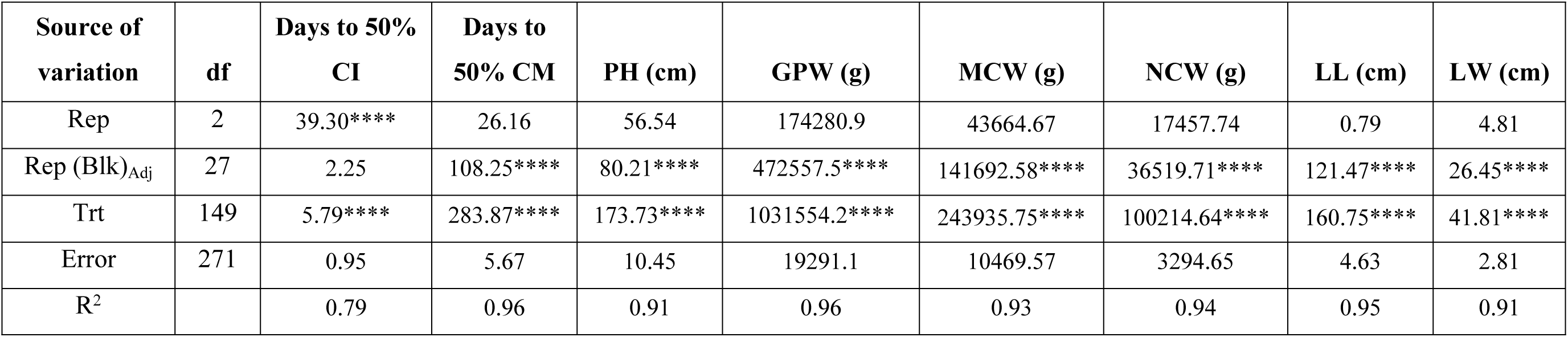
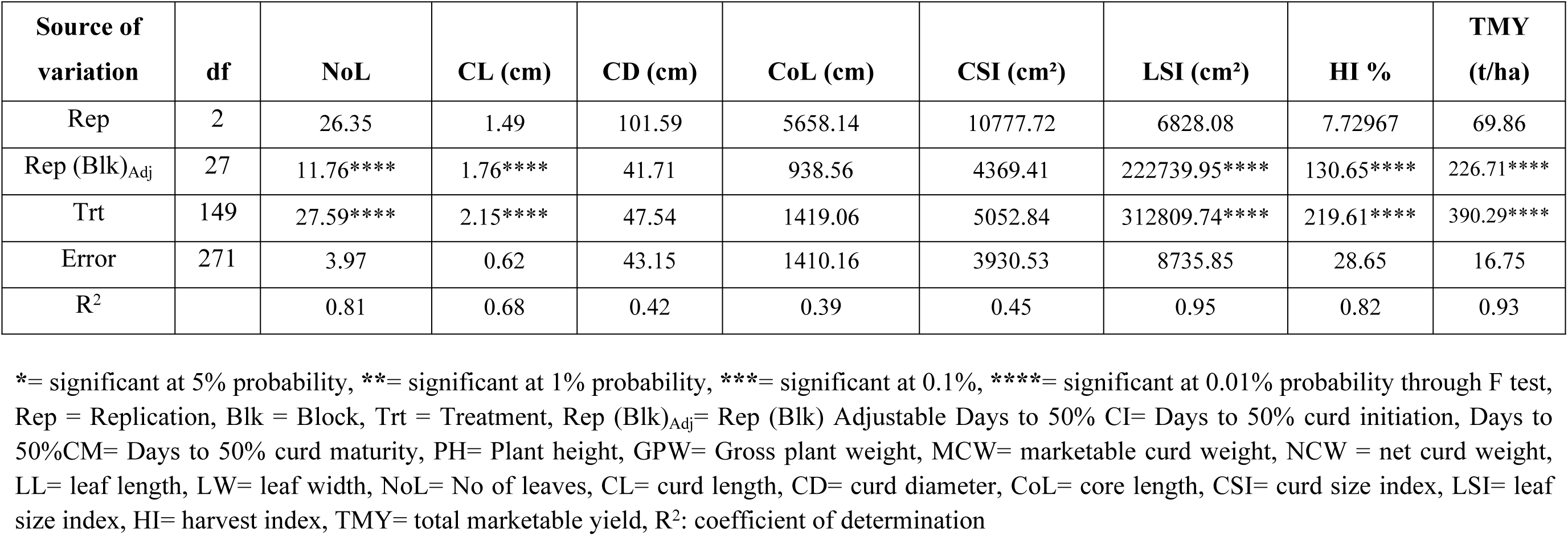
Estimates of Mean Squares and R^2^ for vegetative and commercial traits in Alpha Lattice Design

| Source of variation | df | Days to 50% CI | Days to 50% CM | PH (cm) | GPW (g) | MCW (g) | NCW (g) | LL (cm) | LW (cm) |
| --- | --- | --- | --- | --- | --- | --- | --- | --- | --- |
| Rep | 2 | 39.30**** | 26.16 | 56.54 | 174280.9 | 43664.67 | 17457.74 | 0.79 | 4.81 |
| Rep (Blk)<sub>Adj</sub> | 27 | 2.25 | 108.25**** | 80.21**** | 472557.5**** | 141692.58**** | 36519.71**** | 121.47**** | 26.45**** |
| Trt | 149 | 5.79**** | 283.87**** | 173.73**** | 1031554.2**** | 243935.75**** | 100214.64**** | 160.75**** | 41.81**** |
| Error | 271 | 0.95 | 5.67 | 10.45 | 19291.1 | 10469.57 | 3294.65 | 4.63 | 2.81 |
| R<sup>2</sup> |  | 0.79 | 0.96 | 0.91 | 0.96 | 0.93 | 0.94 | 0.95 | 0.91 |

**Table 2. Continue**
| Source of variation | df | NoL | CL (cm) | CD (cm) | CoL (cm) | CSI (cm<sup>2</sup>) | LSI (cm<sup>2</sup>) | HI % | TMY (t/ha) |
| --- | --- | --- | --- | --- | --- | --- | --- | --- | --- |
| Rep | 2 | 26.35 | 1.49 | 101.59 | 5658.14 | 10777.72 | 6828.08 | 7.72967 | 69.86 |
| Rep (Blk)<sub>Adj</sub> | 27 | 11.76**** | 1.76**** | 41.71 | 938.56 | 4369.41 | 222739.95**** | 130.65**** | 226.71**** |
| Trt | 149 | 27.59**** | 2.15**** | 47.54 | 1419.06 | 5052.84 | 312809.74**** | 219.61**** | 390.29**** |
| Error | 271 | 3.97 | 0.62 | 43.15 | 1410.16 | 3930.53 | 8735.85 | 28.65 | 16.75 |
| R<sup>2</sup> |  | 0.81 | 0.68 | 0.42 | 0.39 | 0.45 | 0.95 | 0.82 | 0.93 |

\*= significant at 5% probability, \*\*= significant at 1% probability, \*\*\*= significant at 0.1%, \*\*\*\*= significant at 0.01% probability through F test, Rep = Replication, Blk = Block, Trt = Treatment, Rep (Blk)<sub>Adj</sub>= Rep (Blk) Adjustable Days to 50% CI= Days to 50% curd initiation, Days to 50%CM= Days to 50% curd maturity, PH= Plant height, GPW= Gross plant weight, MCW= marketable curd weight, NCW = net curd weight, LL= leaf length, LW= leaf width, NoL= No of leaves, CL= curd length, CD= curd diameter, CoL= core length, CSI= curd size index, LSI= leaf size index, HI= harvest index, TMY= total marketable yield, R<sup>2</sup>: coefficient of determination

**Table 3.** Line x Tester Analysis of variance (ANOVA) for combining ability for yield and horticultural traits in cauliflower

| Source of variation | df | Days to 50% CI | Days to 50% CM | PH (cm) | GPW (g) | MCW (g) | NCW (g) | LL (cm) | LW (cm) |
| --- | --- | --- | --- | --- | --- | --- | --- | --- | --- |
| Replicates | 2 | 0.25 | 0.83 | 8.75 | 23153.58 | 11384.77 | 2565.82 | 1.65 | 6.04* |
| Treatments | 145 | 6.08*** | 299.66*** | 176.18*** | 1095365.25*** | 266871.75*** | 106378.93*** | 175.89*** | 43.16*** |
| Parents | 25 | 13.28*** | 563.97*** | 113.34*** | 403231.69*** | 157665.16*** | 56107.28*** | 106.85*** | 29.26*** |
| Parents (Line) | 19 | 15.26*** | 598.47*** | 116.59*** | 324516.00*** | 129477.23*** | 47982.00*** | 104.48*** | 34.83*** |
| Parents (Testers) | 5 | 7.26*** | 35.52*** | 32.52* | 93011.02*** | 92945.55*** | 59482.50*** | 72.38*** | 13.26*** |
| Parents (L vs T) | 1 | 5.75* | 2550.53*** | 455.84*** | 3449932.75*** | 1016833.69*** | 193611.64*** | 324.21*** | 3.48 |
| Parents vs Crosses | 1 | 29.58*** | 1020.72*** | 5878.71*** | 31342492.00*** | 4855010.50*** | 1962991.75*** | 2546.75*** | 330.86*** |
| Crosses | 119 | 4.37*** | 238.08*** | 141.46*** | 986594.00*** | 251258.53*** | 101338.41*** | 170.48*** | 43.66*** |
| Line Effect | 19 | 5.53 | 1054.16*** | 222.35* | 1737689.88* | 335689.69 | 114123.88 | 270.94* | 86.83** |
| Tester Effect | 5 | 5.06 | 33.96 | 65.61 | 694204.25 | 153128.39 | 47422.77 | 180.44 | 49.83 |
| Line * Tester Eff. | 95 | 4.10*** | 85.61*** | 129.27*** | 851763.75*** | 239537.05*** | 101618.98*** | 149.86*** | 34.71*** |
| Error | 290 | 1.20 | 4.94 | 11.04 | 20306.63 | 10387.84 | 3296.66 | 4.67 | 1.96 |
| Total | 437 | 2.82 | 102.71 | 65.82 | 377032.50 | 95495.77 | 37496.81 | 61.47 | 15.65 |

**Table 3 continue**
| Source of variation | df | NoL | CL (cm) | CD (cm) | CoL (cm) | CSI (cm<sup>2</sup>) | LSI (cm<sup>2</sup>) | HI % | TMY (t/ha) |
| --- | --- | --- | --- | --- | --- | --- | --- | --- | --- |
| Replicates | 2 | 2.92 | 1.28 | 1.07 | 0.01 | 181.72 | 3407.57 | 6.81 | 18.22 |
| Treatments | 145 | 29.54*** | 2.35*** | 5.11*** | 1.23*** | 1325.33*** | 348863.53*** | 238.79*** | 426.99*** |
| Parents | 25 | 38.75*** | 1.07* | 3.70*** | 0.95*** | 736.22*** | 169525.92*** | 305.09*** | 252.26*** |
| Parents (Line) | 19 | 40.96*** | 1.21* | 4.14*** | 1.08*** | 820.98*** | 190813.19*** | 323.77*** | 207.16*** |
| Parents (Testers) | 5 | 15.33*** | 0.78 | 2.77*** | 0.61*** | 561.34** | 88306.48*** | 292.54*** | 148.71*** |
| Parents (L vs T) | 1 | 113.78*** | 0.00 | 0.00 | 0.01 | 0.17 | 171165.05*** | 12.92 | 1626.93*** |
| Parents vs Crosses | 1 | 0.79 | 61.33*** | 168.99*** | 16.64*** | 43832.41*** | 3937259.75*** | 1887.12*** | 7768.02*** |
| Crosses | 119 | 27.84*** | 2.12*** | 4.03*** | 1.16*** | 1091.89*** | 356384.91*** | 211.01*** | 402.01*** |
| Line Effect | 19 | 33.27 | 2.20 | 3.84 | 1.39 | 1097.96 | 670563.13** | 226.60 | 537.10 |
| Tester Effect | 5 | 23.30 | 0.10 | 3.10 | 0.58 | 264.71 | 413817.59 | 132.45 | 245.01 |
| Line * Tester Eff. | 95 | 27.00*** | 2.21*** | 4.11*** | 1.15*** | 1134.21*** | 290526.47*** | 212.03*** | 383.26*** |
| Error | 290 | 4.07 | 0.64 | 0.61 | 0.11 | 157.99 | 8677.53 | 27.14 | 16.62 |
| Total | 437 | 12.51 | 1.21 | 2.11 | 0.48 | 545.43 | 121529.77 | 97.27 | 152.79 |

\*, \*\*, \*\*\*, \*\*\*\* significant at 5%, 1%, 0.1%, 0.01% probability respectively through F test,

### Gene action, genetic components of variance, heritability

The estimation of genetic components of variance, nature of gene action, heritability, genetic advance and degree of dominance is presented in Table 4. The GCA variance (σ^2^_gca_) for both lines and testers was found lower in contrast to SCA variance (σ^2^_sca_) for all the vegetative and commercial yield related traits except for Days to 50% CM for which the σ^2^_gca_ for lines was superior than both σ^2^_gca_ for testers and σ^2^_sca_. Then, the value of dominance variance (σ^2^D) was greater as compared to additive component of variance (σ^2^A) for all the studied traits except for Days to 50% CM. The degree of dominance was observed greater than unity for all the studied traits indicating dominant nature of these traits except for Days to 50% CM, for which the value of degree of dominance was approaching to unity (0.99). Further, the ratio of additive to dominance variance (σ^2^A/D) coupled with predictability ratio (σ^2^_gca_/σ^2^_sca_) was found less than unity for all the traits suggesting preponderance of non-additive gene action, except for Days to 50% CM for which the value of σ^2^A/D was slightly higher than unity (1.03). The estimation of heritability magnitude is associated with selection efficiency. In the present investigation, the lowest estimate of narrow-sense heritability (h2ns) was found for CL (3.44%) and highest h^2^_ns_ value was recorded for Days to 50% CM (49.21%). Generally, moderate level of h^2^_ns_ estimates was found for majority of the traits except for CL, CD, CSI and HI, for which low h^2^_ns_ was observed. The higher estimates of genetic advance (GA) at 5% selection intensity were observed for GPW, MCW, NCW and LSI, while lower estimates of GA were recorded for all other traits.

**Table 4.**
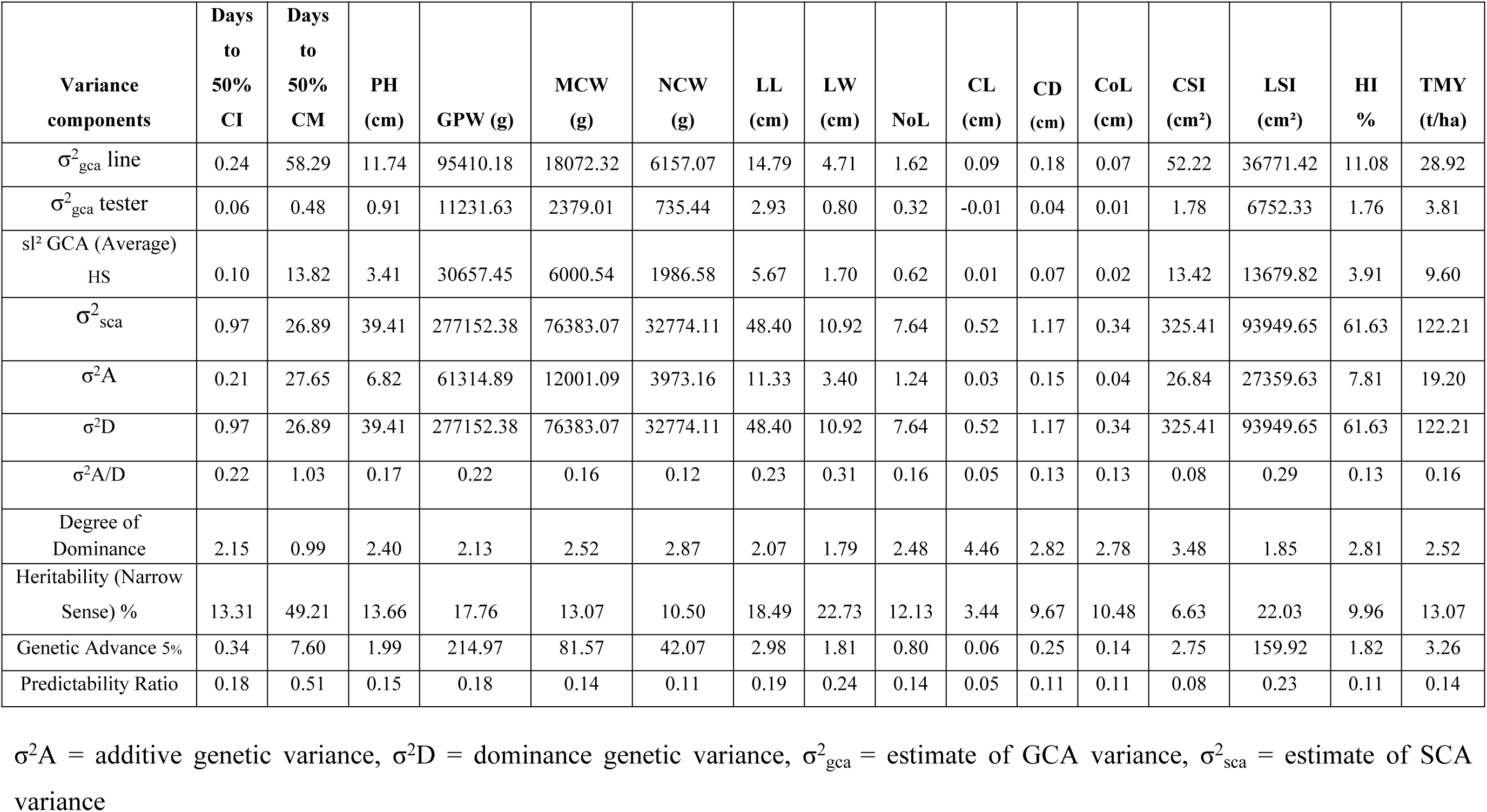
Estimates of genetic components of variance, heritability, genetic advance and predictability ratio for sixteen vegetative and commercial traits

### Combining ability effects

The estimates of combining ability are effective for early generation selection of inbred lines and identifying heterotic crosses. The GCA estimates of parental lines and testers are summarized in Table 5. The GCA estimates revealed that the CMS lines Ogu118-6A, Ogu33A, Ogu34-1A and Ogu33-1A were having significantly high GCA in desirable direction with respect to traits related to earliness such as days to 50% CI and days to 50% CM (Table 5). Besides, the CMS lines Ogu307-33A, Ogu119-1A, Ogu125-8A and tester DH-53-10 also showed significantly high GCA for days to 50% CM in desirable direction (Table 5). The CMS line Ogu13-85-6A was found poor general combiner for all the traits except NCW, CoL and NoL. For the CoL, the significantly high GCA in desirable negative direction was observed in CMS lines Ogu122-5A, Ogu118-6A, Ogu1A, Ogu13-85-6A, Ogu1-6A, Ogu122-1A and tester DH-53-10 (Table 5). For the PH, GPW, MCW and NCW, significantly high GCA in desirable direction was observed in 6, 9, 6 and 9 CMS lines, respectively. While among the six testers, only 2, 1, 2, 2 testers displayed significantly high GCA in desirable direction for these traits respectively. Among the 20 CMS lines used as female parents, 8, 9, 6 and 8 lines showed significantly high GCA in desirable direction for LL, LW, NoL and LSI, respectively. For the curd traits like CL, CD and CSI, 2, 5, 5 CMS lines, respectively, were found good general combiner in positive direction, while 4 CMS lines for each of these traits significantly had negative GCA effects. None of the tester exhibited significant GCA for CL in any direction, while for CD, 1 tester had significantly high GCA in positive direction. For the HI, 7 CMS lines and 1 DH tester (DH-53-9) had significantly high GCA in positive direction. Among the 20 CMS lines, 6 lines (Ogu122-5A, Ogu33A, OguKt-2-6A, Ogu1-6A, Ogu126-1A and Ogu125-8A) had significantly high GCA for TMY in positive direction. While among the six testers, 2 testers, DH-53-1 and DH53-10 exhibited significantly high GCA for TMY in positive direction (Table 5).

**Table 5.** Estimates of general combining ability (GCA) effects of lines and testers

| Lines/testers | Days to<br>50% CI | Days to<br>50% CM | PH (cm) | GPW (g) | MCW (g) | NCW (g) | LL (cm) | LW (cm) |
| --- | --- | --- | --- | --- | --- | --- | --- | --- |
| Ogu122-5A | 0.73** | 6.78*** | -0.10 | 278.71*** | 213.54*** | 31.59* | 0.14 | 1.06** |
| Ogu115-33A | 0.62* | 5.56*** | -2.29** | 330.54*** | 26.21 | 24.64 | 5.09*** | 1.59*** |
| Ogu118-6A | -0.60* | -7.99*** | 0.68 | 274.38*** | 23.93 | 148.70*** | 1.19* | 2.26*** |
| Ogu307-33A | -0.49 | -13.16*** | -0.03 | 209.32*** | 11.21 | -14.02 | 5.17*** | 2.61*** |
| Ogu309-2A | 0.17 | 6.39*** | -5.51*** | -267.13*** | -5.07 | 77.64*** | -4.70*** | -1.80*** |
| Ogu33A | -0.71** | -13.27*** | 0.74 | 402.88*** | 179.27*** | 50.42*** | -1.23* | -0.24 |
| OguKt-2-6A | -0.16 | 4.84*** | 7.55*** | 609.65*** | 223.04*** | 77.20*** | 6.84*** | 5.56*** |
| Ogu1A | -0.27 | 4.73*** | -2.50** | 19.88 | 18.27 | -27.74* | -0.79 | 0.39 |
| Ogu13-85-6A | 0.06 | -0.77 | -3.32*** | 64.04 | -41.46 | 40.31** | -6.09*** | -3.54*** |
| Ogu1-6A | -0.44 | 5.95*** | -0.32 | -40.40 | 62.66** | 66.81*** | -5.78*** | -2.02*** |
| Ogu2A | 0.56* | 4.51*** | -0.84 | -266.40*** | -159.57*** | -68.63*** | -2.00*** | -1.74*** |
| OguKt-9-2A | 0.68** | 8.34*** | 2.87*** | -282.07*** | -135.73*** | -56.63*** | -0.18 | 0.84* |
| Ogu22-1A | 0.12 | 2.95*** | 3.36*** | -109.96** | -39.34 | -53.13*** | -2.38*** | -2.14*** |
| Ogu122-1A | 0.56* | 6.34*** | 3.31*** | -148.29*** | -105.84*** | -84.13*** | 4.09*** | -0.22 |
| Ogu126-1A | 0.56* | 4.01*** | 1.83* | 76.99* | 115.16*** | 67.81*** | 4.55*** | 0.65* |
| Ogu12A | 0.56* | 6.89*** | -0.89 | -459.68*** | -131.51*** | -60.47*** | -3.57*** | -1.59*** |
| Ogu119-1A | -0.05 | -8.05*** | 1.36 | -339.13*** | -206.07*** | -115.13*** | 0.27 | -0.81* |
| Ogu34-1A | -1.10*** | -11.61*** | 1.70* | 78.49* | 25.99 | -0.97 | 1.78*** | 1.05** |
| Ogu125-8A | -0.10 | -2.99*** | 1.42 | 171.71*** | 170.43*** | 71.14*** | 2.30*** | 1.30*** |
| Ogu33-1A | -0.71** | -9.44*** | -9.03*** | -603.51*** | -245.12*** | -175.41*** | -4.70*** | -3.22*** |
| Testers |  |  |  |  |  |  |  |  |
| DH-18-8-1 | 0.25 | 1.29*** | -0.70 | -145.43*** | -67.08*** | -39.01*** | -2.78*** | -1.39*** |
| DH-18-8-3 | 0.16 | 0.28 | -0.59 | -27.38 | -2.74 | -25.86*** | 0.85*** | 0.25 |
| DH-53-1 | -0.40** | -0.36 | 1.38** | 183.02*** | 78.32*** | 31.56*** | 2.12*** | 1.29*** |
| DH-53-6 | -0.34* | -0.72* | 0.86* | -3.38 | -35.79** | -3.36 | -1.09*** | -0.62*** |
| DH-53-9 | 0.11 | 0.14 | -1.35** | -38.20* | -1.61 | 11.89 | 0.95*** | 0.25 |
| DH-53-10 | 0.21 | -0.64* | 0.40 | 31.37 | 28.91* | 24.76*** | -0.06 | 0.22 |
| CD 95%<br>GCA(Line) | 0.51 | 1.03 | 1.54 | 66.17 | 47.32 | 26.66 | 1.00 | 0.65 |
| CD 95%<br>GCA(Tester) | 0.28 | 0.57 | 0.84 | 36.24 | 25.92 | 14.60 | 0.55 | 0.36 |

\*, \*\*, \*\*\*, \*\*\*\* Significance at $P \leq 0.05$ , $P \leq 0.01$ , $P \leq 0.001$ , $P \leq 0.0001$ , respectively, CD: critical difference

**Table 5. Continue**
| Lines/testers | NoL | CL (cm) | CD (cm) | CoL (cm) | CSI (cm<sup>2</sup>) | LSI (cm<sup>2</sup>) | HI % | TMY (t/ha) |
| --- | --- | --- | --- | --- | --- | --- | --- | --- |
| Ogu122-5A | -0.48 | -0.06 | -0.17 | -0.30*** | -2.18 | 42.94 | 3.07* | 8.54*** |
| Ogu115-33A | 0.79 | 0.15 | 0.18 | 0.26** | 2.84 | 198.54*** | -5.79*** | 1.05 |
| Ogu118-6A | 1.63*** | 0.05 | -0.07 | -0.23** | -1.02 | 132.96*** | -5.21*** | 0.96 |
| Ogu307-33A | 0.29 | -0.34 | 0.17 | -0.07 | -3.82 | 261.97*** | -3.71** | 0.45 |
| Ogu309-2A | -1.54** | -0.41* | -0.72*** | -0.05 | -12.58*** | -200.01*** | 6.05*** | -0.20 |
| Ogu33A | 1.13* | 0.33 | 0.01 | 0.00 | 3.38 | -40.80 | -1.78 | 7.17*** |
| OguKt-2-6A | -0.15 | 0.21 | 0.38* | -0.15 | 5.62 | 445.65*** | -4.05** | 8.92*** |
| Ogu1A | 0.96* | 0.31 | 0.48** | -0.48*** | 8.43** | -4.68 | -0.23 | 0.73 |
| Ogu13-85-6A | 1.41** | -0.47* | -0.70*** | -0.22** | -12.56*** | -281.45*** | -1.56 | -1.66 |
| Ogu1-6A | -2.43*** | 0.25 | 0.77*** | -0.26** | 9.57** | -231.80*** | 2.47* | 2.51** |
| Ogu2A | 1.91*** | -0.12 | 0.20 | -0.11 | 0.13 | -143.18*** | -2.02 | -6.38*** |
| OguKt-9-2A | -3.15*** | 0.35 | -0.24 | 0.22** | 2.37 | 28.93 | -0.53 | -5.43*** |
| Ogu22-1A | -1.48** | -0.49* | -0.46* | 0.70*** | -3.55 | -142.26*** | 1.93 | -1.57 |
| Ogu122-1A | -0.59 | -0.09 | -0.16 | -0.22** | -2.54 | 79.39*** | -0.97 | -4.23*** |
| Ogu126-1A | 0.29 | 0.48* | 0.25 | 0.15 | 7.58* | 139.82*** | 3.39** | 4.61*** |
| Ogu12A | -0.65 | -0.06 | -0.22 | 0.23** | -3.34 | -163.41*** | 5.69*** | -5.26*** |
| Ogu119-1A | 0.96* | 0.12 | 0.64*** | 0.26*** | 8.00** | -48.09* | -2.46* | -8.24*** |
| Ogu34-1A | 0.91 | -0.34 | -0.18 | -0.06 | -5.98* | 103.02*** | -1.81 | 1.04 |
| Ogu125-8A | 0.68 | 0.66*** | 0.66*** | 0.36*** | 14.24*** | 116.42*** | 3.52** | 6.82*** |
| Ogu33-1A | -0.48 | -0.54** | -0.81*** | -0.04 | -14.60*** | -293.94*** | 4.00** | -9.80*** |
| Testers |  |  |  |  |  |  |  |  |
| DH-18-8-1 | -0.01 | 0.01 | 0.36*** | 0.17*** | 3.35* | -135.48*** | 1.04 | -2.68*** |
| DH-18-8-3 | -0.98*** | -0.07 | -0.19 | -0.07 | -0.99 | 36.61** | 0.58 | -0.11 |
| DH-53-1 | 0.14 | 0.03 | 0.16 | -0.02 | 1.53 | 100.79*** | -1.71* | 3.13*** |
| DH-53-6 | 0.02 | 0.03 | -0.25* | 0.00 | -2.57 | -54.16*** | -1.87** | -1.43** |
| DH-53-9 | 0.97*** | -0.02 | -0.04 | 0.04 | -0.77 | 39.02** | 1.78** | -0.06 |
| DH-53-10 | -0.14 | 0.03 | -0.05 | -0.12** | -0.56 | 13.21 | 0.19 | 1.16* |
| CD 95%<br>GCA(Line) | 0.94 | 0.37 | 0.36 | 0.16 | 5.84 | 43.25 | 2.42 | 1.89 |
| CD 95%<br>GCA(Tester) | 0.51 | 0.20 | 0.20 | 0.09 | 3.20 | 23.69 | 1.32 | 1.04 |

\*, \*\*, \*\*\*, \*\*\*\* Significance at $P \leq 0.05$ , $P \leq 0.01$ , $P \leq 0.001$ , $P \leq 0.0001$ , respectively, CD: critical difference

The results pertaining to SCA effects of 120 cross combinations are presented in supplementary S2 Table. Among the 120 hybrids, 9 and 28 crosses respectively, showed significantly negative SCA effects for earliness traits, days to 50% CI and days to 50% CM (S2 Table).

The highest SCA effect in desirable negative direction for days to 50% CI was recorded in hybrid Ogu307-33A × DH-18-8-3 (non significant GCA × poor combiner) followed by Ogu34-1A × DH-53-1(good general combiner × good combiner) and Ogu309-2A × DH-53-9 (poor combiner × poor combiner). For the days to 50% CM, the highest SCA effect in desirable negative direction was observed in the cross Ogu22-1A × DH-53-10 (poor combiner × good general combiner) followed by Ogu125-8A × DH-53-6 (good general combiner × good combiner) and Ogu1A × DH-18-8-3 (poor combiner × poor combiner). Among the 120 testcross progenies, 31 crosses exhibited high significant positive SCA effects for PH (S2 Table). The highest positive significant SCA effect for PH was observed in the cross Ogu122-1A × DH-18-8-3 (good general combiner × poor combiner) followed by Ogu13-85-6A × DH-53-1 (poor combiner × good combiner), Ogu13-85-6A × DH-18-8-3 (poor combiner × poor combiner) and Ogu118-6A × DH-53-9 (non significant GCA × poor combiner). For the commercial traits viz. GPW, MCW, NCW, CL, CD, CoL and CSI, out of 120 crosses, 39, 32, 38, 10, 26, 33 and 24 crosses exhibited significantly high SCA effects in desirable direction (S2 Table). For the vegetative traits, among 120 crosses, 44, 37, 33 and 48 crosses showed significantly high positive SCA effects for LL, LW, NoL and LSI, respectively. For the HI and TMY, among 120 hybrids, 18 and 32 hybrids displayed significantly high positive SCA effects, respectively (S2 Table). The cross combination Ogu22-1A × DH-53-6 (poor combiner × poor combiner) exhibited highest positive significant SCA effect for GPW followed by Ogu307-33A × DH-18-8-3 (good general combiner × poor combiner) and Ogu122-5A × DH-53-10 (good general combiner × non significant GCA). For the MCW, hybrid Ogu33A × DH53-1 (good combiner × good combiner) showed highest positive significant SCA effects followed by Ogu122-5A × DH-53-10 (good combiner × good combiner) and Ogu1-6A × DH-53-1 (good combiner × good combiner). The highest significantly positive SCA estimate for NCW was observed in the cross Ogu33A × DH-53-1 (good combiner × good combiner) followed by Ogu1A × DH-53-9 (poor combiner × non significant GCA) and Ogu22-1A × DH-53-6 (poor combiner × poor combiner). With respect to CL, the highest significant positive SCA effect was observed in the cross Ogu119-1A × DH18-8-1 (non significant GCA × non significant GCA) followed by Ogu122-1A × DH-53-10 (poor combiner × non significant GCA) and Ogu122-5A × DH-53-10 (poor combiner × non significant GCA), likewise for CD, the highest positive SCA effect was recorded in the cross combination Ogu13-85-6A × DH-18-8-3 (poor combiner × poor combiner) followed by Ogu2A × DH-53-6 (non significant GCA × poor combiner) and Ogu122-1A × DH-53-10 (poor combiner × poor combiner). Then, for the CoL the highest significant SCA effect in desirable negative direction was observed in the cross Ogu12A × DH-53-10 (poor combiner × good general combiner) followed by Ogu2A × DH-18-8-1 (non significant GCA × poor combiner) and Ogu119-1A × DH-53-1 (poor combiner × non significant GCA). The cross combination Ogu119-1A × DH-18-8-1 (good combiner × good combiner) exhibited highest positive SCA effect for CSI. The crosses Ogu13-85-6A × DH-53-1 (poor combiner × good combiner) followed by Ogu118-6A × DH-53-9 (good combiner × good combiner) and Ogu12A × DH-18-8-1 (poor combiner × poor combiner) displayed highest significant positive SCA effect for LL, similarly for LW, the highest positive significant SCA estimate was observed in the hybrid Ogu12A × DH-18-8-1 (poor combiner × poor combiner) followed by Ogu33-1A × DH-53-9 (poor combiner × non significant GCA) and Ogu22-1A × DH-53-1 (poor combiner × good combiner). With respect to NoL, the highest positive significant SCA effect was recorded in cross combination OguKt2-6A × DH-53-10 (poor combiner × poor combiner) followed by Ogu2A × DH-53-9 (good combiner × good combiner) and Ogu13-85-6A × DH-18-8-1 (good combiner × poor combiner). For the LSI, the cross Ogu12A × DH-18-8-1 (poor combiner × poor combiner) exhibited highest significant positive SCA effect. Then, the crosses Ogu33A × DH-53-1 (poor combiner × poor combiner) followed by Ogu125-8A × DH-18-8-1 (good combiner × non significant GCA) and Ogu122-5A × DH-53-9 (good combiner × good combiner) displayed highest significant positive SCA effects for HI. For the total marketable yield (TMY), the highest significant SCA estimate in desirable positive direction was observed in the cross Ogu33A × DH-53-1 (good combiner × good combiner) followed by Ogu122-5A × DH-53-10 (good combiner × good combiner) and Ogu1-6A × DH-53-1 (good combiner × good combiner) (S2 Table).

### Mean performance and cluster analysis based on agronomic traits

The mean performance of parental CMS and DH lines along with standard checks is presented in supplement S3 Table. On the basis of curd initiation, the CMS lines Ogu307-33A and Ogu13-85-6A were earliest among rest of parental lines and the entire four standard checks as well (S3 Table). Similarly, CMS lines Ogu307-33A and Ogu33-1A were earliest among all the parents and checks with respect to curd maturity. Then CMS line Ogu22-1A, OguKt-9-2A, Ogu12A and DH lines DH-53-1, DH-53-10 had highest plant height. While, CMS lines Ogu115-33A and Ogu309-2A was having dwarfed structure in contrast to other parental lines. The highest number of leaves was observed in CMS lines Ogu34-1A and Ogu309-2A. The shortest core length was recorded in genotype Ogu13-85-6A and Ogu309-2A. The tester DH-53-1 was having highest MCW. The highest total marketable yield was recorded in CMS line Ogu33A and Ogu125-8A, whereas the tester DH-18-8-1 was having highest TMY among all the parental genotypes and standard checks. Principal component analysis (PCA) and hierarchical cluster analysis (HCA) based on 16 agronomic traits (days to 50% CI, days to 50% CM, PH, GPW, MCW, NCW, LL, LW, NoL, CL, CD, CoL, LSI, CSI, HI and TMY) were performed for grouping of 26 parental CMS and DH inbred lines of 120 testcross progenies (Figs 1a and b). The PCA revealed that, the first two dimensions (PC1 and PC2) captured 36.2% and 18.9% of total existing variation among the parental lines. The HCA based dendrogram depicting inter-relationships displayed high genetic divergence among 26 CMS and DH lines based upon Euclidean distance matrix (Fig 1b). The HCA of 26 parental lines on the basis of 16 phenotypic traits classified parental CMS and DH lines into 3 major clusters with varying extent of divergence within internal sub-clusters. The DH lines DH-53-10 and DH-18-8-1 was distantly placed from rest of DH testers in two different major clusters.

**Fig1a.**
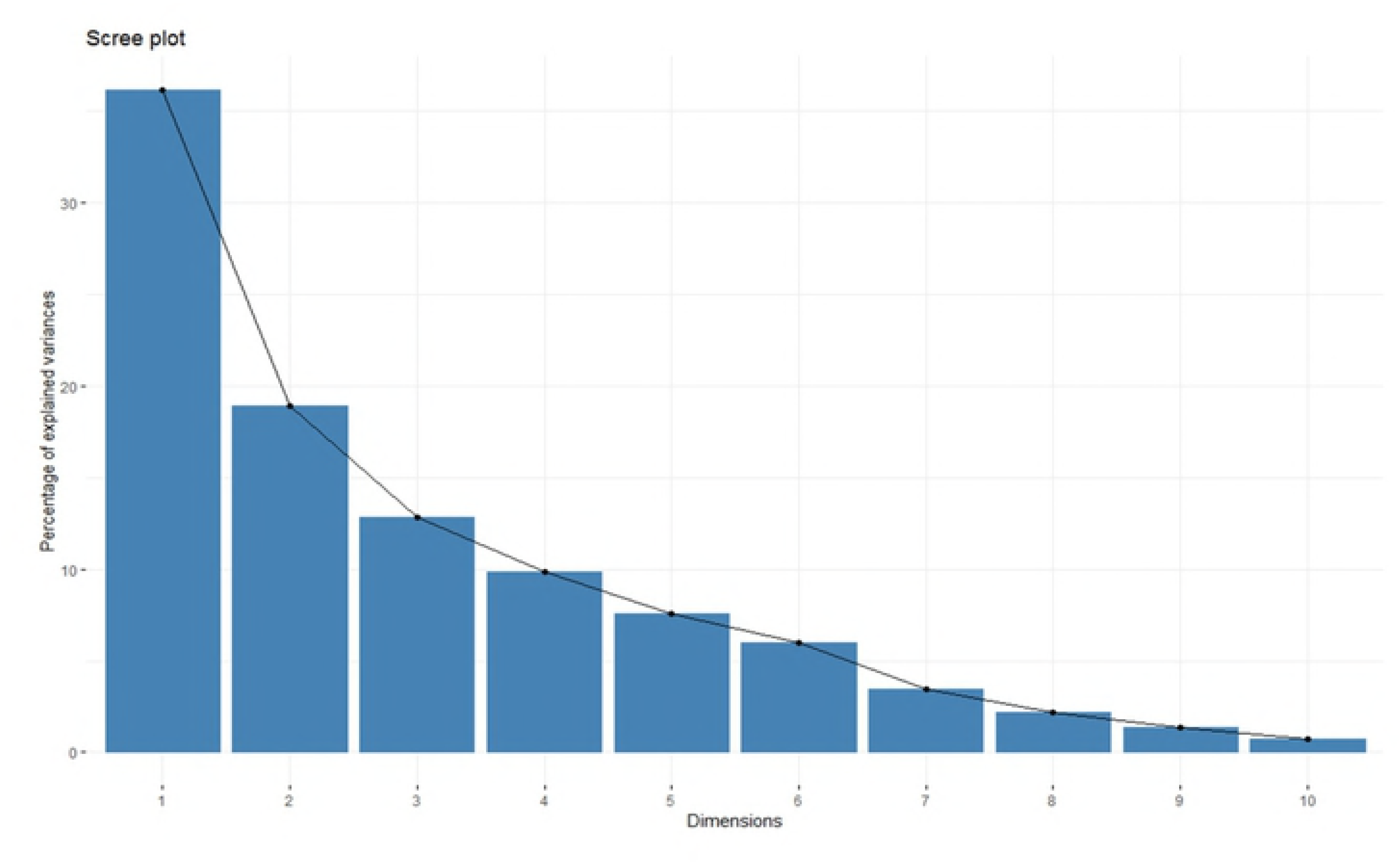
Percentage of explained variance among parental CMS and DH lines in different principal components

**Fig1b.**
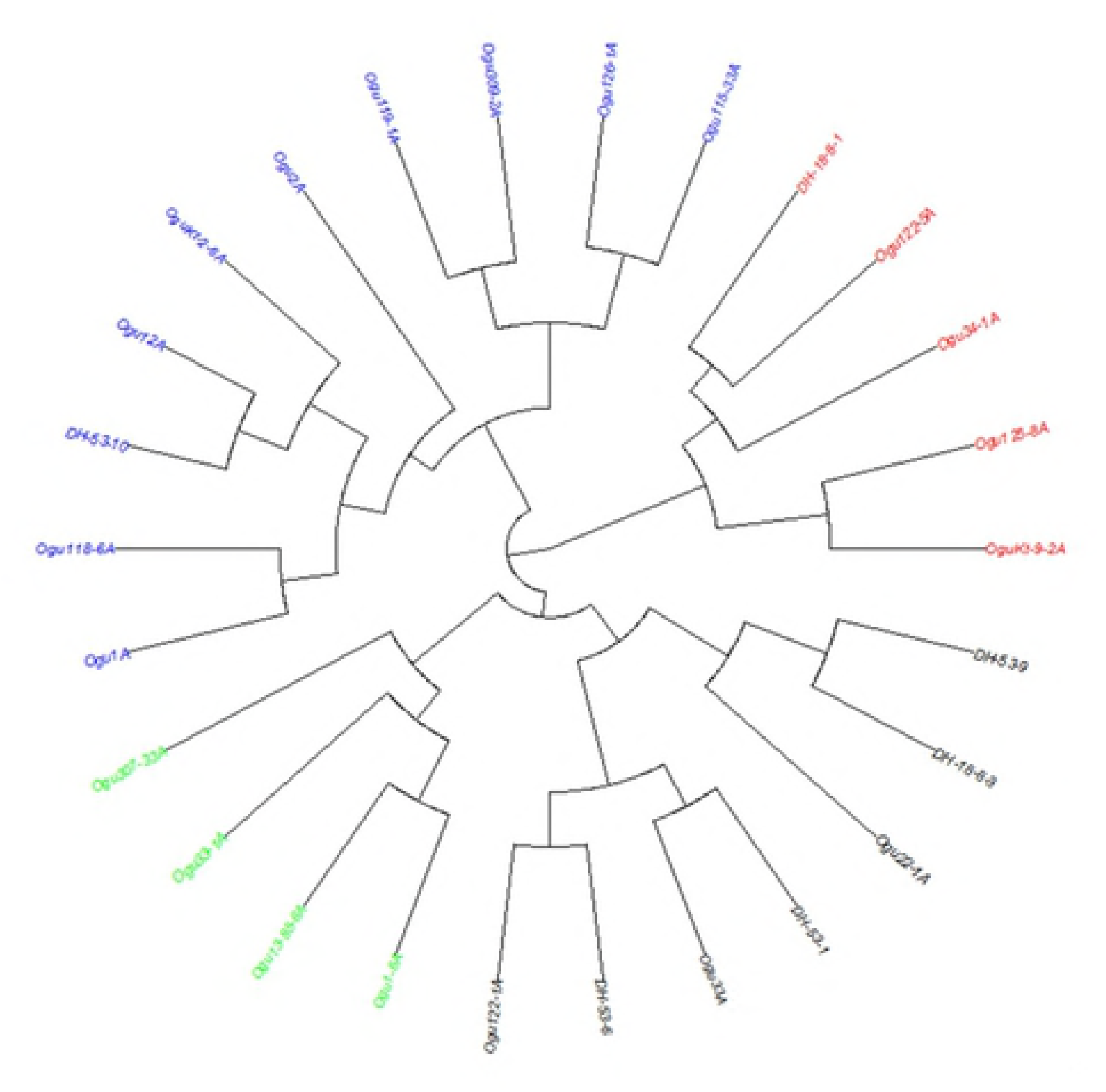
Dendrogram illustrating the genetic relationships among 26 parental lines based on 16 phenotypic traits.

### SSR and EST-SSRs based polymorphism, allelic diversity and genetic distances

In the present investigation, 350 pairs of microsatellite markers (genomic-SSR and EST-SSRs) based primers distributed throughout the *Brassica oleracea* genome were tested to assess the molecular diversity in parental CMS and DH lines of cauliflower. Out of 350 microsatellite primers, 87 pairs of primers displayed clear cut polymorphism and revealed high allelic diversity (Table 6). The allele frequency analysis revealed that overall 511 alleles were amplified through 87 microsatellite primers (Table 6) with mean number of alleles per locus was 5.87. The allele numbers per locus ranged from 2 (1 primer BoSF1640) to 10 (1 primer: BRAS011) (Table 6). The observed heterozygosity (*H_o_*) ranged from 0.03 (for the loci BoSF2232, BoSF062, BRAS011, BoESSR080, BoSF2406 and BoSF2421) to 0.19 (for the loci BoSF2294a). The mean expected heterozygosity (*H_e_*) was 0.68, with a range of 0.27 (primer cnu107) to 0.83 (for primer Na12F03a and BoESSR041) and had higher mean value than *H_o_*. The mean polymorphic information content (PIC) for 87 loci was 0.63. The PIC content ranged from 0.24 for the primer cnu107 to 0.80 for the primer Na12F03a (Table 6). Further, the PCA and Neighbour joining (NJ) cluster analysis based on molecular data for 87 loci, revealed distinct clusters and sub-clusters of parental CMS and DH lines based on their phylogeny (Fig 2). The PCA revealed that first two major coordinate axis 1 and 2 (PC1 and PC2) explained 61.41% of total existing variation among CMS and DH lines. The dendrogram constructed revealed 3 main clusters of parental lines with internal sub-clusters showing varying degree of diversity. As the less extent of variation explained by first two main coordinate axes (PC1 and PC2), the NJ clusters gave clear picture of clustering groups for better interpretation. The DH testers remained in 2 different sub-clusters of single main cluster. The CMS lines Ogu2A and OguKt-9-2A placed distantly from rest of CMS lines. The CMS lines Ogu33-1A and Ogu125-8A were in close affinity with DH lines.

**Table 6.** Characteristics of 87 polymorphic SSR and EST-SSRs loci depicting diversity

| Locus | LG | H<sub>o</sub> | H<sub>e</sub> | N | PIC | Locus | LG | H<sub>o</sub> | H<sub>e</sub> | N | PIC |
| --- | --- | --- | --- | --- | --- | --- | --- | --- | --- | --- | --- |
| BoSF2304b | C09 | 0.00 | 0.48 | 5 | 0.43 | BoESSR303 | C04 | 0.00 | 0.33 | 4 | 0.31 |
| BoSF1740 | C08 | 0.00 | 0.63 | 5 | 0.57 | BoESSR333 | C04 | 0.00 | 0.81 | 9 | 0.77 |
| BoSF378 | C08 | 0.00 | 0.72 | 4 | 0.65 | BoESSR338 | C08 | 0.00 | 0.77 | 8 | 0.72 |
| BoSF2680 | C08 | 0.00 | 0.78 | 5 | 0.73 | BoESSR403 | C08 | 0.00 | 0.72 | 7 | 0.66 |
| BoSF2054 | C06 | 0.00 | 0.52 | 3 | 0.41 | BoESSR409 | C04 | 0.00 | 0.78 | 6 | 0.73 |
| BoSF1215 | C06 | 0.00 | 0.59 | 6 | 0.54 | BoESSR576 | C06 | 0.00 | 0.71 | 6 | 0.65 |
| BoSF250 | C06 | 0.00 | 0.67 | 3 | 0.58 | BoESSR581 | C06 | 0.00 | 0.67 | 7 | 0.62 |
| BoSF2505 | C06 | 0.00 | 0.60 | 4 | 0.54 | BoESSR632 | C01 | 0.00 | 0.71 | 6 | 0.67 |
| BoSF2374 | C05 | 0.00 | 0.62 | 4 | 0.54 | BoESSR901 | C09 | 0.00 | 0.77 | 7 | 0.73 |
| BoSF1846 | C05 | 0.00 | 0.69 | 4 | 0.62 | BoESSR766 | C03 | 0.00 | 0.69 | 7 | 0.63 |
| BoSF2878 | C05 | 0.00 | 0.57 | 3 | 0.48 | BoESSR825 |  | 0.00 | 0.69 | 7 | 0.63 |
| BoSF912 | C01 | 0.00 | 0.72 | 6 | 0.67 | BoESSR673 | C03 | 0.00 | 0.66 | 5 | 0.58 |
| BoSF063 | C01 | 0.00 | 0.74 | 8 | 0.69 | BoESSR758 | C07 | 0.00 | 0.68 | 7 | 0.64 |
| BoSF2294a | C02 | 0.19 | 0.73 | 6 | 0.67 | BoESSR763 | C04 | 0.00 | 0.80 | 8 | 0.76 |
| BoSF2615 | C02 | 0.00 | 0.61 | 4 | 0.55 | BoESSR863 | C06 | 0.00 | 0.69 | 4 | 0.62 |
| BoSF1167 | C02 | 0.00 | 0.79 | 6 | 0.75 | BoESSR903 | C06 | 0.00 | 0.50 | 5 | 0.46 |
| BoSF2248 | C03 | 0.00 | 0.60 | 5 | 0.55 | Na12F03a | C07 | 0.07 | 0.83 | 8 | 0.80 |
| BoSF2232 | C03 | 0.03 | 0.49 | 3 | 0.38 | O110B11 | C05 | 0.00 | 0.65 | 4 | 0.57 |
| BoSF184 | C04 | 0.00 | 0.73 | 7 | 0.68 | BoSF2406 | C07 | 0.03 | 0.67 | 7 | 0.62 |
| BoSF1640 | C09 | 0.00 | 0.43 | 2 | 0.33 | BoSF2313 | C07 | 0.07 | 0.75 | 8 | 0.70 |
| BoSF2612 | C08 | 0.00 | 0.67 | 4 | 0.59 | BoSF2033 | C07 | 0.00 | 0.81 | 9 | 0.77 |
| BoSF2860 | C07 | 0.04 | 0.76 | 8 | 0.71 | BoSF317 | C05 | 0.00 | 0.70 | 6 | 0.66 |
| BoSF2345 | C01 | 0.00 | 0.76 | 5 | 0.71 | BoSF2421 | C09 | 0.03 | 0.61 | 6 | 0.57 |
| BoSF1207 | C01 | 0.07 | 0.77 | 8 | 0.73 | BoSF1957 | C04 | 0.00 | 0.82 | 8 | 0.77 |
| BoSF042 | C03 | 0.00 | 0.73 | 6 | 0.68 | Na12B09 | C03 | 0.00 | 0.70 | 6 | 0.66 |
| BoSF062 | C03 | 0.03 | 0.51 | 5 | 0.46 | cnu107 | C02 | 0.00 | 0.27 | 3 | 0.24 |
| BoSF2985 | C03 | 0.00 | 0.68 | 4 | 0.60 | BoSF1131 | C03 | 0.00 | 0.78 | 7 | 0.73 |
| BoE862 | C04 | 0.00 | 0.75 | 6 | 0.70 | BoSF966 | C03 | 0.00 | 0.80 | 7 | 0.75 |
| BRAS011 | C02 | 0.03 | 0.77 | 10 | 0.73 | CB10258 | C01 | 0.00 | 0.78 | 8 | 0.73 |
| BrBAC214 | C03 | 0.00 | 0.48 | 5 | 0.43 | BoESSR920 | C09 | 0.00 | 0.79 | 6 | 0.73 |
| BoESSR080 | C07 | 0.03 | 0.78 | 9 | 0.73 | BoESSR041 | C06 | 0.00 | 0.83 | 8 | 0.79 |
| BoESSR086 | C03 | 0.00 | 0.63 | 6 | 0.58 | BoESSR934 | C08 | 0.00 | 0.65 | 4 | 0.57 |
| BoESSR087 | C04 | 0.00 | 0.70 | 5 | 0.65 | Ni4D12 | C02 | 0.00 | 0.57 | 3 | 0.48 |
| BoESSR089 | C01 | 0.00 | 0.80 | 7 | 0.76 | cnu149 | C05 | 0.00 | 0.81 | 8 | 0.76 |
| BoESSR105 | C04 | 0.00 | 0.76 | 6 | 0.71 | BoESSR482 | C02 | 0.00 | 0.79 | 7 | 0.75 |
| BoESSR108 | C04 | 0.00 | 0.67 | 5 | 0.60 | O112G04a | C08 | 0.00 | 0.44 | 4 | 0.40 |
| BoESSR122 | C02 | 0.00 | 0.82 | 8 | 0.78 | BoESSR492 | C03 | 0.00 | 0.76 | 8 | 0.72 |
| BoESSR151 | C02 | 0.00 | 0.70 | 6 | 0.65 | BoESSR510 | C03 | 0.00 | 0.59 | 4 | 0.51 |
| BoESSR206 | C05 | 0.00 | 0.53 | 4 | 0.47 | BoESSR523 | C07 | 0.00 | 0.76 | 7 | 0.71 |
| BoESSR207 | C05 | 0.00 | 0.78 | 5 | 0.72 | BoESSR560 | C03 | 0.00 | 0.73 | 6 | 0.67 |
| BoESSR208 | C04 | 0.00 | 0.71 | 8 | 0.66 | BoESSR736 | C05 | 0.00 | 0.66 | 5 | 0.61 |
| BoESSR212 | C07 | 0.00 | 0.56 | 4 | 0.51 | BoESSR030 | C03 | 0.00 | 0.76 | 8 | 0.71 |
| BoESSR216 | C01 | 0.00 | 0.76 | 5 | 0.70 | BoESSR073 | C03 | 0.00 | 0.50 | 6 | 0.46 |
| BoESSR248 | C04 | 0.00 | 0.62 | 5 | 0.57 |  |  |  |  |  |  |

LG: linkage group, H<sub>o</sub>: observed heterozygosity, H<sub>e</sub>: expected heterozygosity, PIC: polymorphic information content

**Fig 2.**
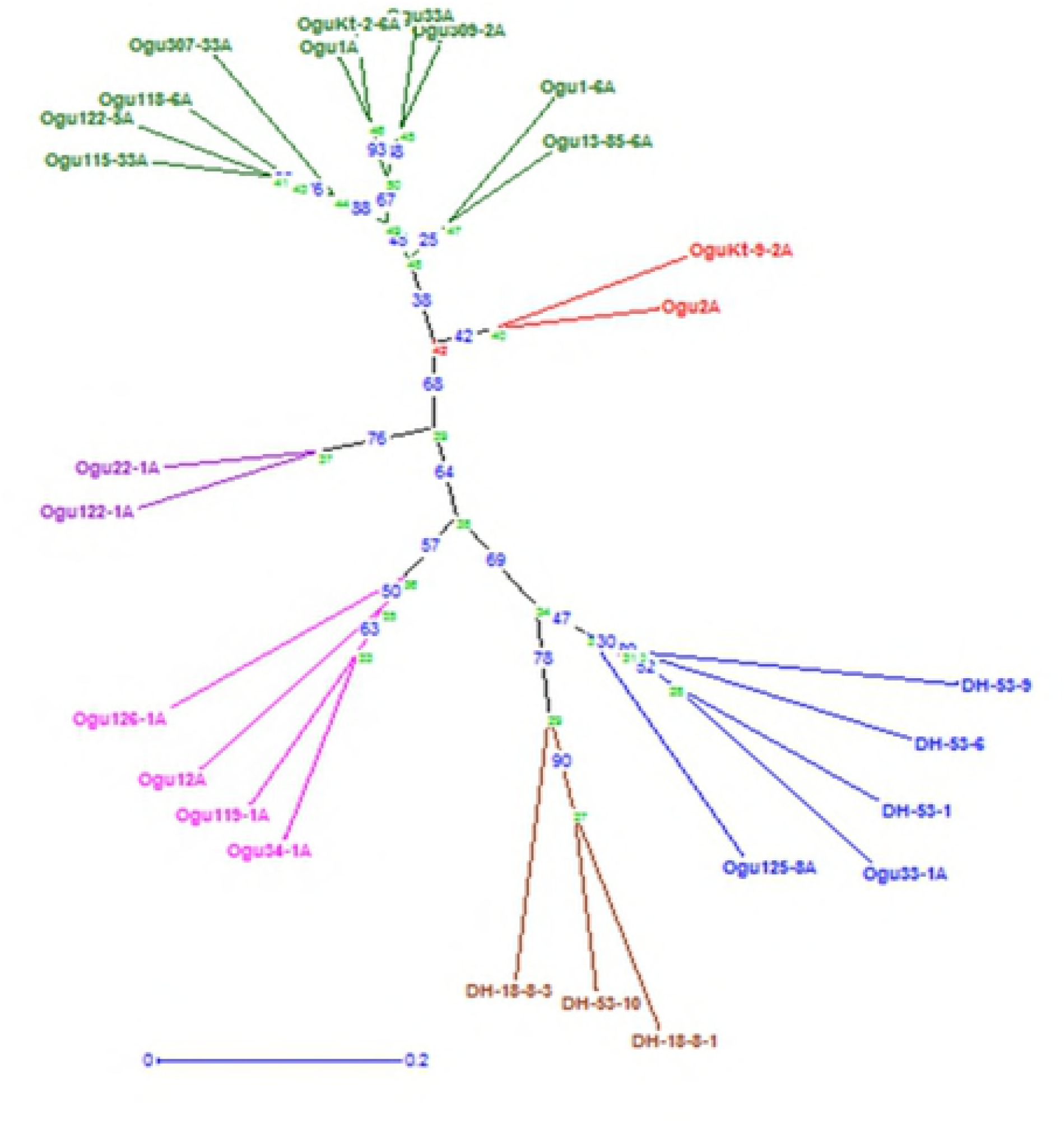
Dendrogram of parental lines through UPGMA cluster analysis illustrating the genetic relationships among them based on SSR and EST-SSR analysis (molecular data).

The Euclidean distance (PD) between lines and testers were computed from 16 phenotypic traits (supplement Table S3) and GD was calculated from molecular data based on 87 microsatellite markers (genomic-SSR and EST-SSRs) used for assessment of genetic diversity between parents (S4 Table). The PD was ranged from 2.07 for the cross L16 × T6 (Ogu12A × DH-53-10) to 8.27 for the cross combination L5 × T1 (Ogu309-2A × DH-18-8-1) with a mean of 5.52. The GD was ranged from 0.44 for the cross L20 × T1 (Ogu33-1A × DH-18-8-1) to 0.98 for the cross combinations L4 × T5 (Ogu307-33A × DH-53-9) with the average GD of 0.83.

### Genetic structure analysis

To infer pedigree and genetic clusters of 26 parental inbred lines, genetic structure analysis was performed using Bayesian approach by STRUCTURE version 2.3.4 under admixture model with correlated allele frequencies and as far as possible this model attempts to identify population clusters which are not in disequilibrium. The range of demes (*k*) tested was *k* =1 to *k* = 10 with 15 runs for each *k* to quantify the extent of variation of the likelihood for each *k.* The result of analysis by STRUCTURE HARVESTER version v0.6.94 revealed that second order likelihood, *ΔK* reached to peak at *k* = 4 (Figs 3a to 3c), hence, optimal *k* value should be 4. This indicated that 26 parental CMS and DH inbred lines could be grouped into 4 genetic sub-clusters (CI: first cluster depicted by red color, CII: second cluster represented by light green color, CIII: third cluster is represented by blue color and CIV: fourth sub-cluster by yellow color) (Fig 3c). All the DH testers were remained in same cluster III depicted in blue color, including 2 CMS lines Ogu125-8A and Ogu33-1A, which remained in the vicinity of DH testers. Although there is minor admixture in DH-53-10 and Ogu125-8A from the genotypes of cluster I and cluster II respectively, indicating somewhat gene flow in the cluster III from cluster I and cluster II. The other CMS lines placed themselves in separate clusters. Thus 20 CMS lines used as female parent of 120 testcross progenies were grouped into 4 sub-clusters. The maximum number of CMS lines were placed in cluster I depicted by red color. There is admixture from cluster IV to the cluster I and cluster II genotypes Ogu1A, Ogu13-85-6A and Ogu122-1A, respectively. Similarly, there was minor admixture from cluster I and cluster II to cluster IV genotypes (Ogu1-6A and Ogu22-1A). Thus, there were four distinct four sub-clusters including minor gene flow within some genotypes of respective clusters from each other.

**Fig 3.**
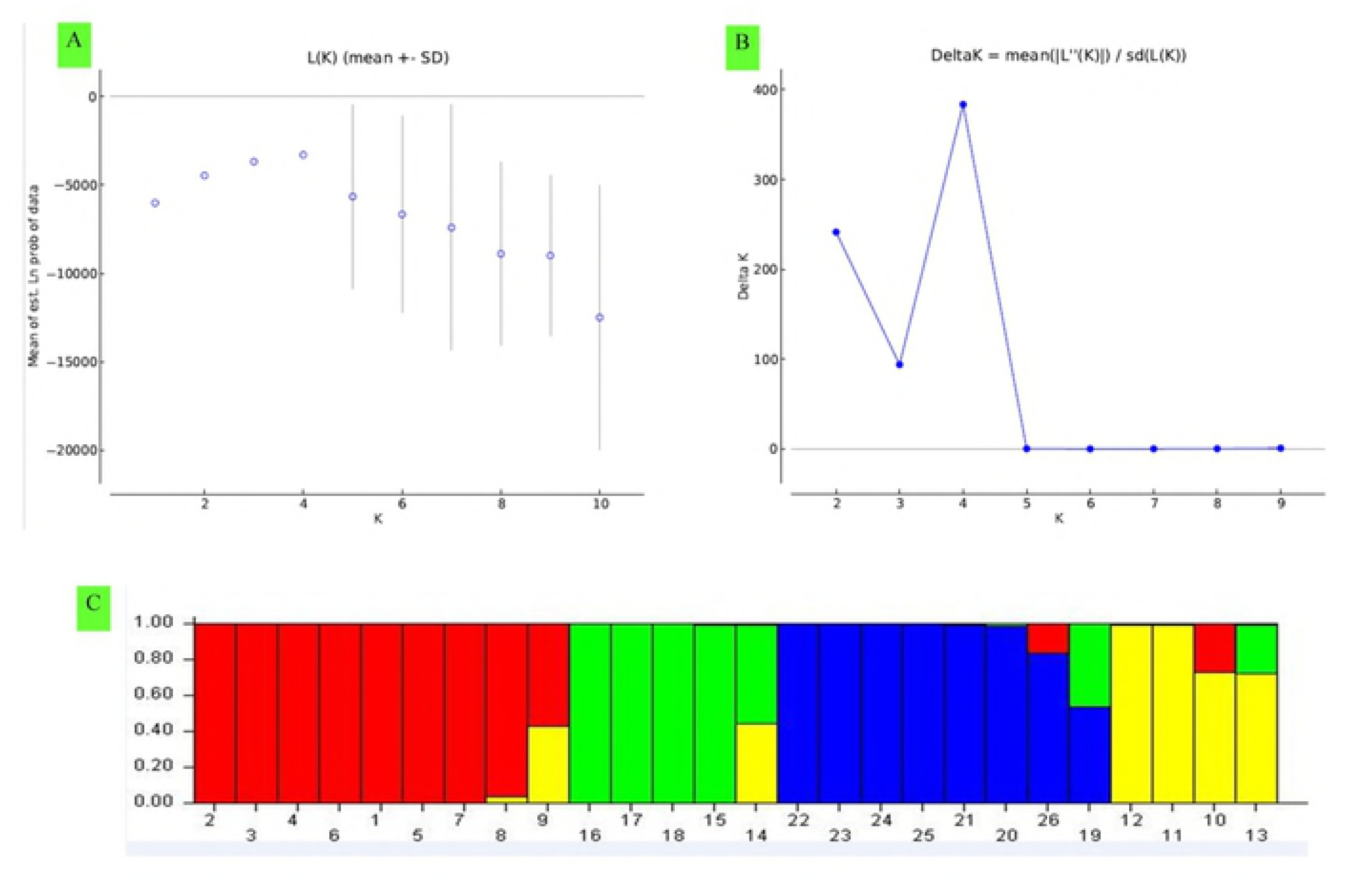
Genetic structure analysis of parental CMS and DH lines by STRUCTURE v2.3.4 and STRUCTURE HARVESTER based upon 87 SSR, EST-SSR loci. (a) Mean L(k) ± SD over 15 runs for each *k* value from 1 to 10. (b) *ΔK calculated as ΔK = m*|*L*′′ (*K*)|/*s* [*L*(*K*)], reached peak at *k* = 4. (c) Q-plot clustering. Inferred ancestries of CMS and DH lines based on 4 genetic groups. Each cluster is represented by different color and each column represent respective genotype allotted to respective cluster. Different color of each column depicts the percent of membership (vertical values on the left of cluster) of each genotype for four clusters.

### Analysis of heterosis

The heterotic response of all the 120 testcross progenies varied in magnitude and highly significant heterosis (MPH, BPH) was observed for all the 16 traits in both directions (data not presented). The top ten cross combinations based on significant MPH in desirable direction along with their BPH and SCA effects, for all the 16 vegetative and commercial traits, respectively, are presented in Table 7 and Table 8. Among the vegetative traits, for the traits related with earliness, like days to 50% CI and days to 50% CM, the cross combinations Ogu34-1A×DH-53-1 and Ogu33A×DH-53-6, showed significantly high MPH in desirable negative direction (Table 7). For the days to 50% CI, the testers DH-53-1 and DH-53-9, were involved in 4 crosses individually out of top 10 crosses. For the days to 50% curd maturity, the CMS line Ogu33A as female parent was involved in 6 hybrids for earliness among top 10 hybrids. Ogu33A was also involved as female parent in one of the top 10 cross combinations related to days to 50% CI. This line had significantly highest GCA for earliness among all the CMS lines used in the study. Thus, CMS line Ogu33A could be used as good parent for generating early F_1_ hybrids in cauliflower. For the PH, among the top 10 heterotic crosses, the cross combination Ogu118-6A×DH-53-9 exhibited highest significant positive heterosis over mid-parent followed by OguKt-2-6A×DH-53-9 and Ogu34-1A×DH-53-9. The highest significant positive heterosis for GPW was observed in the cross Ogu118-6A×DH-53-10 over mid-parent followed by Ogu126-1A×DH-53-1 and Ogu307-33A×DH-18-8-3. The highest significant MPH for NoL in desirable direction was found in the cross combination OguKt-2-6A×DH-53-10 followed by Ogu115-33A×DH-53-10 and Ogu1A×DH-53-6, likewise with respect to LSI, the highest positive significant MPH was observed in the cross Ogu126-1A×DH-53-1 followed by OguKt-2-6A×DH-53-1 and Ogu126-1A×DH-53-10. With respect to LL, the cross Ogu126-1A×DH-53-1 showed highest significant MPH in positive direction followed by OguKt-2-6A×DH-53-1 and Ogu115-33A×DH-18-8-3. For the LW, the cross combination OguKt-2-6A×DH-53-1 exhibited highest significant positive heterosis over mid-parent followed by Ogu126-1A×DH-53-10 and Ogu126-1A×DH-53-1. The cross OguKt-2-6A×DH-53-1 was highest and second highest among top 10 crosses, with significant positive MPH for LW and LL, respectively. The CMS lines OguKt-2-6A had significantly high positive GCA for both LL and LW. The top ten crosses having significant positive MPH for 8 commercial traits are presented in Table 8.

**Table 7.** MPH of top ten crosses along with their BPH, mean performance and SCA effects (value in parenthesis) for 8 vegetative traits

| Days to 50% curd initiation |  |  |  | Days to 50% curd maturity |  |  |  |
| --- | --- | --- | --- | --- | --- | --- | --- |
| Cross combination | MPH% | BPH% | Mean performance | Cross combination | MPH% | BPH% | Mean performance |
| Ogu34-1A×DH-53-1 | -6.35** (-3.26***) | -7.19** | 86.00 | Ogu33A×DH-53-6 | -17.11** (-4.94***) | -17.11** | 126.00 |
| Ogu2A×DH-53-1 | -5.43** (-0.93) | -7.85** | 90.00 | Ogu33A×DH-53-1 | -15.81** (-2.64*) | -16.27** | 128.66 |
| Ogu2A×DH-53-9 | -5.37** (-0.45) | -6.83** | 91.00 | Ogu33A×DH-53-9 | -13.41** (-0.48) | -13.60** | 131.33 |
| Ogu33A×DH-53-1 | -4.69** (-1.65**) | -5.04** | 88.00 | Ogu33A×DH-18-8-1 | -12.58** (-3.29*) | -14.69** | 129.66 |
| Ogu309-2A×DH-53-9 | -4.50** (-2.72***) | -6.69** | 88.33 | Ogu22-1A×DH-53-10 | -12.54** (-13.58***) | -12.64** | 133.66 |
| Ogu125-8A×DH-53-1 | -3.99** (-1.93**) | -4.68** | 88.33 | Ogu1A×DH-18-8-3 | -10.27** (-11.61***) | -10.37** | 138.33 |
| Ogu307-33A×DH-18-8-3 | -3.59** (-5.44***) | -6.59** | 85.00 | Ogu33A×DH-18-8-3 | -10.24** (5.39***) | -10.82** | 137.33 |
| OguKt-2-6A×DH-53-9 | -3.57** (-0.73) | -4.93** | 90.00 | Ogu33A×DH-53-10 | -10.07** (5.97***) | -10.26** | 137.00 |
| Ogu1A×DH-53-9 | -3.57** (-0.61) | -4.93** | 90.00 | Ogu118-6A×DH-53-6 | -9.24** (-6.89***) | -14.91** | 129.33 |
| Ogu2A×DH-18-8-1 | -3.36** (-0.58) | -6.83** | 91.00 | Ogu2A×DH-18-8-3 | -8.52** (-10.06***) | -9.31** | 139.66 |
| Plant Height (PH) |  |  |  | Gross Plant Weight (GPW) |  |  |  |
| Cross combination | MPH% | BPH% | Mean Performance | Cross combination | MPH% | BPH% | Mean Performance |
| Ogu118-6A×DH-53-9 | 57.77** (11.18***) | 48.05** | 65.63 | Ogu118-6A×DH-53-10 | 170.14** (1018.24***) | 123.75** | 3442.00 |
| OguKt-2-6A×DH-53-9 | 54.21** (6.47***) | 52.93** | 67.80 | Ogu126-1A×DH-53-1 | 122.47** (800.31***) | 76.74** | 3178.33 |
| Ogu34-1A×DH-53-9 | 53.53** (8.96***) | 45.34** | 64.43 | Ogu307-33A×DH-18-8-3 | 121.64** (1172.05***) | 84.71** | 3472.00 |
| Ogu13-85-6A×DH-53-1 | 48.12** (13.61***) | 25.41** | 66.80 | Ogu115-33A×DH-53-9 | 118.03** (692.97***) | 60.66** | 3103.33 |
| OguKt-2-6A×DH-18-8-1 | 45.77** (5.59**) | 37.61** | 67.56 | OguKt-2-6A×DH-53-1 | 108.76** (569.98***) | 93.55** | 3480.66 |
| Ogu13-85-6A×DH-18-8-3 | 45.06** (11.18***) | 27.09** | 62.40 | Ogu307-33A×DH-18-8-1 | 107.02** (834.10***) | 81.65** | 3016.00 |
| Ogu307-33A×DH-18-8-3 | 44.98** (10.93***) | 33.27** | 65.43 | Ogu1-6A×DH-53-1 | 105.85** (849.37***) | 72.94** | 3110.00 |
| Ogu2A×DH-18-8-1 | 44.85** (9.28***) | 28.04** | 62.86 | Ogu22-1A×DH-53-6 | 104.61** (1175.32***) | 58.34** | 3180.00 |
| Ogu34-1A×DH-53-1 | 44.72** (8.99***) | 26.16** | 67.20 | Ogu1A×DH-53-9 | 99.19** (1022.31***) | 61.62** | 3122.00 |
| Ogu115-33A×DH-53-6 | 41.93** (7.58***) | 20.13** | 61.26 | Ogu115-33A×DH-53-1 | 98.53** (61.76) | 49.77** | 2693.33 |

\*= significant at 5% probability, \*\*= significant at 1% probability, \*\*\*= significant at 0.1%, \*\*\*\*= significant at 0.01% probability through F test, MPH: Mid parent heterosis, BPH: better parent heterosis, SCA: specific combining ability (value in parenthesis)

**Table 7. continue**
| Number of leaves (NoL) |  |  |  | Leaf size index (LSI) |  |  |  |
| --- | --- | --- | --- | --- | --- | --- | --- |
| Cross combination | MPH% | BPH% | Mean performance | Cross combination | MPH% | BPH% | Mean Performance |
| OguKt-2-6A×DH-53-10 | 54.35** (5.53***) | 42.00** | 23.66 | Ogu126-1A×DH-53-1 | 155.50** (313.65***) | 129.09** | 1673.66 |
| Ogu115-33A×DH-53-10 | 51.65** (3.92***) | 40.82** | 23.00 | OguKt-2-6A×DH-53-1 | 144.80** (326.37***) | 122.09** | 1992.21 |
| Ogu1A×DH-53-6 | 42.27** (3.59**) | 23.21** | 23.00 | Ogu126-1A×DH-53-10 | 130.23** (397.38***) | 91.70** | 1669.81 |
| Ogu2A×DH-53-9 | 41.59** (5.36***) | 37.93** | 26.66 | Ogu115-33A×DH-53-6 | 118.91** (345.14***) | 169.04** | 1608.92 |
| Ogu118-6A×DH-53-10 | 41.05** (2.42*) | 26.42** | 22.33 | Ogu118-6A×DH-53-1 | 112.80** (70.98) | 94.93** | 1424.14 |
| Ogu33A×DH-53-1 | 40.00** (3.64**) | 25.00** | 23.33 | Ogu115-33A×DH-53-10 | 111.73** (139.55*) | 68.84** | 1470.71 |
| Ogu2A×DH-18-8-3 | 39.39** (3.64**) | 25.45** | 23.00 | Ogu307-33A×DH-53-10 | 99.43** (291.06***) | 93.52** | 1685.65 |
| Ogu12A×DH-53-10 | 37.93** (2.37*) | 33.33** | 20.00 | Ogu34-1A×DH-53-10 | 92.48** (524.64***) | 83.75** | 1760.28 |
| Ogu1A×DH-53-9 | 37.37** (2.31*) | 17.24** | 22.66 | Ogu34-1A×DH-53-1 | 90.16** (282.31***) | 67.59** | 1605.53 |
| Ogu122-1A×DH-53-10 | 35.48** (-2.36**) | 28.57** | 15.33 | Ogu118-6A×DH-53-10 | 89.85** (138.28*) | 61.17** | 1403.87 |
| Leaf Length (LL) |  |  |  | Leaf width (LW) |  |  |  |
| Cross combination | MPH% | BPH% | Mean Performance | Cross combination | MPH% | BPH% | Mean Performance |
| Ogu126-1A×DH-53-1 | 63.74** (4.77***) | 54.68** | 60.06 | OguKt-2-6A×DH-53-1 | 58.71** (2.47**) | 49.30** | 31.90 |
| OguKt-2-6A×DH-53-1 | 54.77** (4.92***) | 49.05** | 62.50 | Ogu126-1A×DH-53-10 | 56.87** (4.90***) | 46.47** | 28.36 |
| Ogu115-33A×DH-18-8-3 | 54.42** (8.01***) | 26.65** | 62.56 | Ogu126-1A×DH-53-1 | 56.22** (3.31***) | 47.79** | 27.83 |
| Ogu115-33A×DH-53-10 | 51.09** (4.11**) | 28.85** | 57.76 | Ogu118-6A×DH-53-1 | 56.21** (1.33) | 45.84** | 27.46 |
| Ogu13-85-6A×DH-53-1 | 49.38** (13.58***) | 48.81** | 58.23 | OguKt-2-6A×DH-53-9 | 53.15** (5.27***) | 48.97** | 33.66 |
| Ogu126-1A×DH-53-10 | 48.26** (5.72***) | 31.23** | 58.83 | Ogu115-33A×DH-53-6 | 53.03** (3.39***) | 43.26** | 26.93 |
| Ogu115-33A×DH-53-9 | 47.97** (6.00***) | 20.45** | 60.66 | Ogu34-1A×DH-53-10 | 47.39** (5.28***) | 44.46** | 29.13 |
| Ogu115-33A×DH-53-6 | 45.18** (7.14***) | 17.88** | 59.76 | Ogu118-6A×DH-53-10 | 46.97** (1.16) | 35.46** | 26.23 |
| Ogu125-8A×DH-53-1 | 40.58** (9.03***) | 25.47** | 62.06 | Ogu118-6A×DH-53-6 | 43.64** (1.01) | 34.22** | 25.23 |
| Ogu118-6A×DH-53-9 | 40.39** (10.71***) | 22.04** | 61.46 | Ogu307-33A×DH-53-10 | 42.47** (2.48**) | 40.91** | 27.90 |

\*= significant at 5% probability, \*\*= significant at 1% probability, \*\*\*= significant at 0.1%, \*\*\*\*= significant at 0.01% probability through F test, MPH: Mid parent heterosis, BPH: better parent heterosis, SCA: specific combining ability (value in parenthesis)

**Table 8.** MPH of top ten crosses along with their better parent heterosis and SCA effects (value in parenthesis) for 8 commercial traits

| Core length (CoL) |  |  |  | Curd length (CL) |  |  |  |
| --- | --- | --- | --- | --- | --- | --- | --- |
| Cross combination | MPH% | BPH% | Mean Performance | Cross combination | MPH% | BPH% | Mean Performance |
| Ogu122-1A×DH-53-6 | -30.31** (-0.88***) | -37.69** | 3.3 | Ogu119-1A×DH-18-8-1 | 36.24** (2.09***) | 25.69** | 11.41 |
| Ogu1A×DH-53-6 | -22.91** (-0.52**) | -27.00** | 3.4 | Ogu126-1A×DH-53-1 | 34.41** (0.75) | 32.56** | 10.45 |
| Ogu1A×DH-53-10 | -22.50** (-0.30) | -25.16** | 3.5 | Ogu122-1A×DH-53-10 | 33.09** (1.69***) | 30.91** | 10.82 |
| Ogu1A×DH-53-1 | -20.61** (-0.46*) | -26.36** | 3.4 | Ogu119-1A×DH-53-6 | 31.29** (0.77) | 30.67** | 10.12 |
| Ogu12A×DH-53-10 | -20.18** (-1.30***) | -26.02** | 3.2 | Ogu115-33A×DH-53-9 | 31.02** (0.74) | 27.97** | 10.06 |
| Ogu2A×DH-53-10 | -13.62** (-0.88***) | -24.20** | 3.3 | Ogu33A×DH-53-1 | 30.29** (0.45) | 26.85** | 10.00 |
| Ogu122-1A×DH-18-8-3 | -11.58** (0.13) | -21.87** | 4.2 | Ogu33A×DH-53-9 | 29.13** (0.39) | 25.85** | 9.90 |
| Ogu34-1A×DH-18-8-3 | -10.83** (-0.82***) | -23.09** | 3.4 | Ogu126-1A×DH-53-6 | 29.08** (0.24) | 28.39** | 9.95 |
| Ogu126-1A×DH-53-10 | -10.53** (-0.91***) | -19.12** | 3.5 | Ogu119-1A×DH-53-10 | 28.55** (0.90) | 23.94** | 10.25 |
| Ogu122-5A×DH-53-6 | 12.16* (0.62**) | 11.28 | 4.7 | Ogu115-33A×DH-53-6 | 27.65** (0.35) | 25.59** | 9.73 |
| Curd diameter (CD) |  |  |  | Curd Size index (CSI) |  |  |  |
| Cross combination | MPH% | BPH% | Mean Performance | Cross combination | MPH% | BPH% | Mean Performance |
| Ogu1-6A×DH-53-1 | 42.58** (1.40**) | 40.80** | 14.81 | Ogu122-1A×DH-53-10 | 84.33** (41.50***) | 73.67** | 154.33 |
| Ogu122-1A×DH-53-10 | 38.30** (1.96***) | 32.40** | 14.23 | Ogu126-1A×DH-53-1 | 80.34** (19.58**) | 78.43** | 144.61 |
| Ogu22-1A×DH-53-6 | 36.73** (1.67***) | 35.69** | 13.43 | Ogu119-1A×DH-18-8-1 | 73.32** (42.09***) | 48.23** | 169.36 |
| Ogu307-33A×DH-18-8-1 | 34.76** (1.72***) | 16.93** | 14.73 | Ogu1-6A×DH-53-1 | 73.01** (17.53*) | 67.98** | 144.55 |
| Ogu126-1A×DH-53-1 | 34.56** (0.97*) | 33.98** | 13.86 | Ogu33A×DH-53-1 | 72.36** (12.40) | 64.40** | 133.24 |
| Ogu115-33A×DH-53-6 | 34.46** (0.35) | 34.34** | 13.30 | Ogu119-1A×DH-53-10 | 72.05** (22.91**) | 64.60** | 146.27 |
| Ogu119-1A×DH-53-10 | 33.81** (1.21**) | 32.78** | 14.27 | Ogu115-33A×DH-53-6 | 70.76** (12.65) | 67.73** | 128.84 |
| Ogu2A×DH-53-6 | 33.71** (2.09***) | 22.84** | 14.52 | Ogu115-33A×DH-53-9 | 69.91** (14.55*) | 61.78** | 132.55 |
| Ogu1-6A×DH-53-6 | 33.51** (0.63) | 29.55** | 13.63 | Ogu119-1A×DH-53-6 | 68.05** (11.40) | 63.54** | 132.75 |
| OguKt-2-6A×DH-53-9 | 32.44** (1.13*) | 30.99** | 13.95 | Ogu2A×DH-53-6 | 66.85** (34.85***) | 46.88** | 148.33 |

\*= significant at 5% probability, \*\*= significant at 1% probability, \*\*\*= significant at 0.1%, \*\*\*\*= significant at 0.01% probability through F test, MPH: Mid parent heterosis, BPH: better parent heterosis, SCA: specific combining ability (value in parenthesis)

**Table 8. continue**
| Marketable curd weight (MCW) |  |  |  | Net curd weight (NCW) |  |  |  |
| --- | --- | --- | --- | --- | --- | --- | --- |
| Cross combination | MPH% | BPH% | Mean Performance | Cross combination | MPH% | BPH% | Mean Performance |
| Ogu126-1A×DH-18-8-3 | 172.31** (676.08***) | 103.25** | 1876.66 | Ogu118-6A×DH-53-10 | 193.70** (332.68***) | 171.33** | 1157.66 |
| Ogu122-5A×DH-53-10 | 130.78** (731.04***) | 100.49** | 2061.66 | Ogu1A×DH-53-9 | 145.53** (415.99***) | 103.55** | 1051.66 |
| Ogu307-33A×DH-18-8-3 | 115.93** (584.02***) | 82.02** | 1680.66 | Ogu119-1A×DH-53-10 | 141.03** (222.18***) | 116.59** | 783.33 |
| Ogu119-1A×DH-53-10 | 104.24** (292.32***) | 58.68** | 1203.33 | Ogu126-1A×DH-18-8-3 | 139.29** (209.86***) | 94.27** | 903.33 |
| Ogu309-2A×DH-53-6 | 102.53** (527.68***) | 42.28** | 1575.00 | Ogu309-2A×DH-53-10 | 132.70** (0.41) | 108.57** | 754.33 |
| Ogu118-6A×DH-53-10 | 99.69** (344.98***) | 95.96** | 1486.00 | Ogu309-2A×DH-53-6 | 118.87** (337.52***) | 55.23** | 1063.33 |
| Ogu307-33A×DH-53-10 | 96.36** (238.04***) | 80.18** | 1366.33 | Ogu33-1A×DH-53-10 | 105.78** (270.79***) | 98.71** | 771.66 |
| Ogu1A×DH-53-9 | 93.86** (525.17***) | 71.88** | 1630.00 | Ogu309-2A×DH-18-8-3 | 101.86** (55.36) | 63.15** | 758.66 |
| Ogu309-2A×DH-53-10 | 93.81** (57.32) | 54.20** | 1169.33 | Ogu122-5A×DH-53-10 | 101.77** (280.79***) | 59.89** | 988.66 |
| Ogu33A×DH-53-1 | 92.76** (906.90***) | 82.90** | 2252.66 | Ogu13-85-6A×DH-18-8-3 | 101.67** (131.61***) | 90.82** | 887.33 |
| Harvest index (HI) |  |  |  | Total marketable yield (TMY) |  |  |  |
| Cross combination | MPH% | BPH% | Mean Performance | Cross combination | MPH% | BPH% | Mean Performance |
| Ogu122-5A×DH-53-9 | 56.64** (17.39***) | 50.91** | 74.56 | Ogu126-1A×DH-18-8-3 | 172.31** (27.04***) | 103.25** | 75.06 |
| Ogu126-1A×DH-18-8-3 | 49.16** (12.52***) | 39.61** | 68.80 | Ogu122-5A×DH-53-10 | 130.78** (29.24***) | 100.49** | 82.46 |
| Ogu12A×DH-18-8-3 | 41.99** (12.71***) | 39.41** | 71.29 | Ogu307-33A×DH-18-8-3 | 115.93** (23.36***) | 82.02** | 67.22 |
| Ogu119-1A×DH-53-10 | 35.84** (10.63***) | 23.25** | 60.69 | Ogu119-1A×DH-53-10 | 104.24** (11.69***) | 58.68** | 48.13 |
| Ogu126-1A×DH-53-10 | 29.08** (3.61) | 20.86** | 59.51 | Ogu309-2A×DH-53-6 | 102.53** (21.11***) | 42.28** | 63.00 |
| Ogu122-5A×DH-53-10 | 23.63** (3.16) | 19.30** | 58.74 | Ogu118-6A×DH-53-10 | 99.69** (13.80***) | 95.96** | 59.44 |
| Ogu122-1A×DH-53-10 | 22.52** (17.08***) | 9.31 | 68.62 | Ogu307-33A×DH-53-10 | 96.36** (9.52***) | 80.18** | 54.65 |
| Ogu33A×DH-53-1 | 21.79** (33.46***) | 19.90** | 82.29 | Ogu1A×DH-53-9 | 93.86** (21.01***) | 71.88** | 65.20 |
| Ogu309-2A×DH-53-6 | 21.64** (9.07**) | 18.47** | 65.57 | Ogu309-2A×DH-53-10 | 93.81** (2.29) | 54.20** | 46.77 |
| Ogu22-1A×DH-53-9 | 21.16** (12.81***) | 7.17 | 68.84 | Ogu33A×DH-53-1 | 92.76** (36.28***) | 82.90** | 90.10 |

\*= significant at 5% probability, \*\*= significant at 1% probability, \*\*\*= significant at 0.1%, \*\*\*\*= significant at 0.01% probability through F test, MPH: Mid parent heterosis, BPH: better parent heterosis, SCA: specific combining ability (value in parenthesis)

A short core length is desirable in cauliflower. The hybrid Ogu122-1A×DH-53-6 exhibited significantly highest MPH for CoL in desirable negative direction followed by Ogu1A×DH-53-6 and Ogu1A×DH-53-10. The CMS line Ogu1A was involved in 3 crosses as female parent among top 4 crosses with respect to CoL, and it had significantly highest GCA in desirable negative direction for CoL. Thus, Ogu1A could be used as parent for developing hybrids with short core.

For the commercial traits CL and CD, the cross combinations Ogu119-1A×DH-18-8-1 followed by Ogu126-1A×DH-53-1 and Ogu122-1A×DH-53-10 for CL, Ogu1-6A×DH-53-1 followed by Ogu122-1A×DH-53-10 and Ogu22-1A×DH-53-6 for CD, were having significantly highest MPH in desirable positive direction. The cross Ogu122-1A×DH-53-10, showed highest significant positive heterosis over mid-parent for CSI. For economic trait MCW, the hybrid Ogu126-1A×DH-18-8-3 followed by Ogu122-5A×DH-53-10 and Ogu307-33A×DH-18-8-3, exhibited significantly highest positive MPH at P ≤ 0.001. Likewise, for the NCW, the significant highest heterosis over mid parent in desirable positive direction was observed in the hybrid Ogu118-6A×DH-53-10 followed by Ogu1A×DH-53-9 and Ogu119-1A×DH-53-10. The cross combination Ogu122-5A×DH-53-9 (56.64%) showed highest significant MPH for percent HI followed by Ogu126-1A×DH-18-8-3 (49.16%) and Ogu12A×DH-18-8-3 (41.99%). For the total marketable yield (TMY), the highest significant heterosis over mid-parent in desirable positive direction was observed in the testcross Ogu126-1A×DH-18-8-3 followed by Ogu122-5A×DH-53-10 and Ogu307-33A×DH-18-8-3. Among the top ten crosses also wide range of MPH was recorded for TMY from 92.76% (Ogu33A×DH-53-1) to 172.31% (Ogu126-1A×DH-18-8-3).

### Association of genetic distances, heterosis and combining ability

The PPMCC of GD, PD with MPH, BPH, SCA and of combining ability with heterosis for ten commercial traits is presented in Table 9 (Figs 4 a to f). The GD and PD exhibited no significant correlation coefficient with SCA for any of the traits (Table 9). SCA showed significantly positive correlation with MPH and BPH for all the traits at P ≤ 0.01. No significant association of GD with MPH and BPH was observed with respect to days to 50% CM, LL, CL and CoL. For the commercial traits viz. PH, GPW, NCW, LW, CD and TMY, significant correlation was observed between GD and MPH in desirable direction for the respective traits (Table 9). The highest value of PPMCC of GD with MPH and BPH in desirable direction was observed for LW. Thus, GD exhibited significant correlation with MPH and BPH for six traits out of ten traits studied. However, PD exhibited no significant correlation with heterosis for majority of traits. PD showed significant correlation with MPH for only LL in desirable direction (Table 9). PD exhibited significant correlation in undesirable direction for CoL. However, no significant association was observed between parental genetic distances based on phenotypic traits (PD) and molecular data (GD) (r = −0.04) (Fig 5).

**Table 9.**
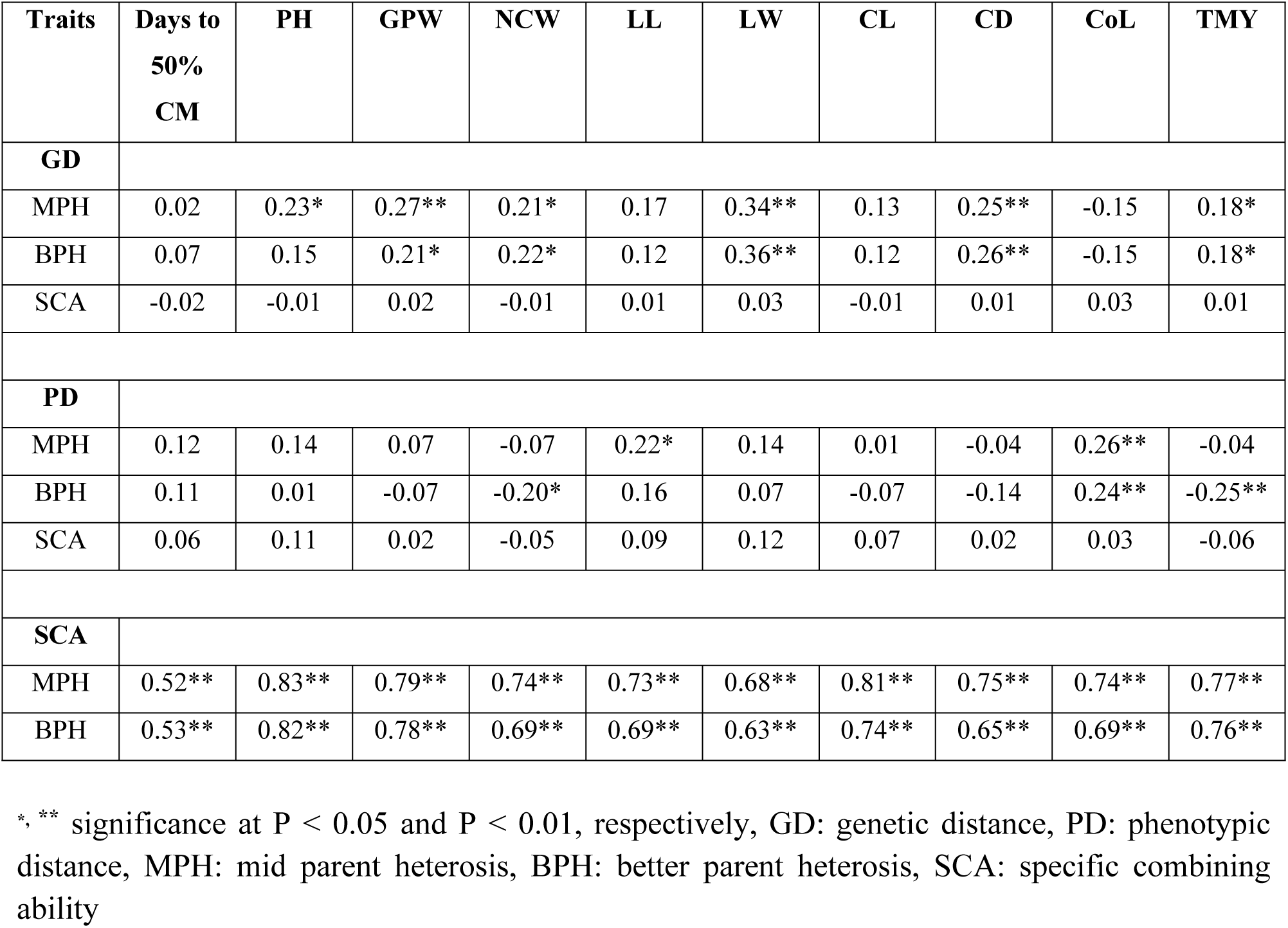
Pearson correlation coefficients among parental genetic distance (GD, PD), combining ability and heterosis in cauliflower for ten morphological and commercial traits

**Fig 4.**
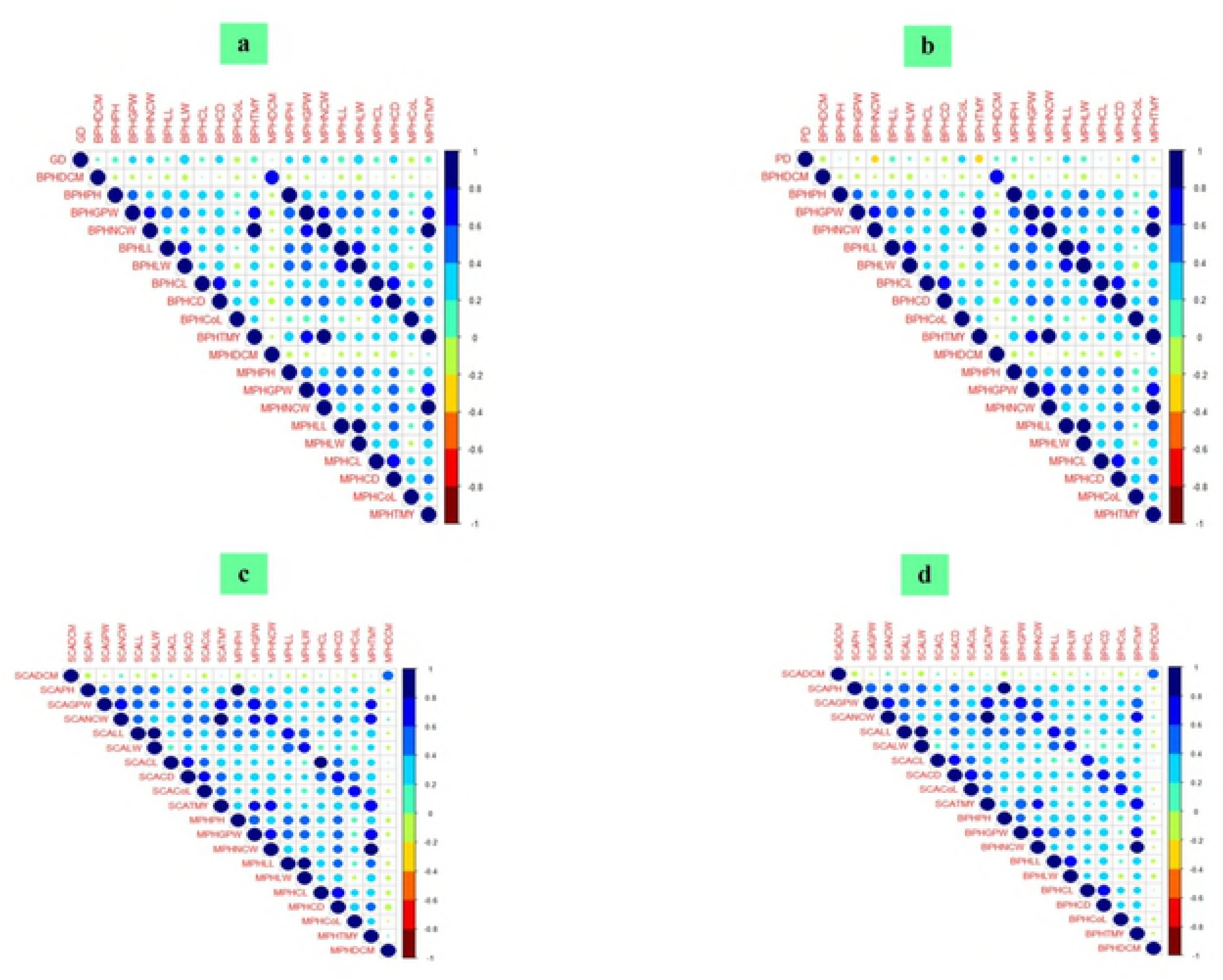
(a, b, c, d) Pearson’s correlation matrix of PD based on phenotypic traits, GD based on SSR, EST-SSR molecular data of 87 loci with MPH, BPH, SCA and SCA with MPH and BPH. a) PPMCC of GD with BPH and MPH for 10 commercial traits, b) PPMCC of PD with MPH and BPH for 10 commercial traits, c) SCA with MPH for 10 commercial traits, d) SCA with BPH for 10 commercial traits.

**Fig.4.**
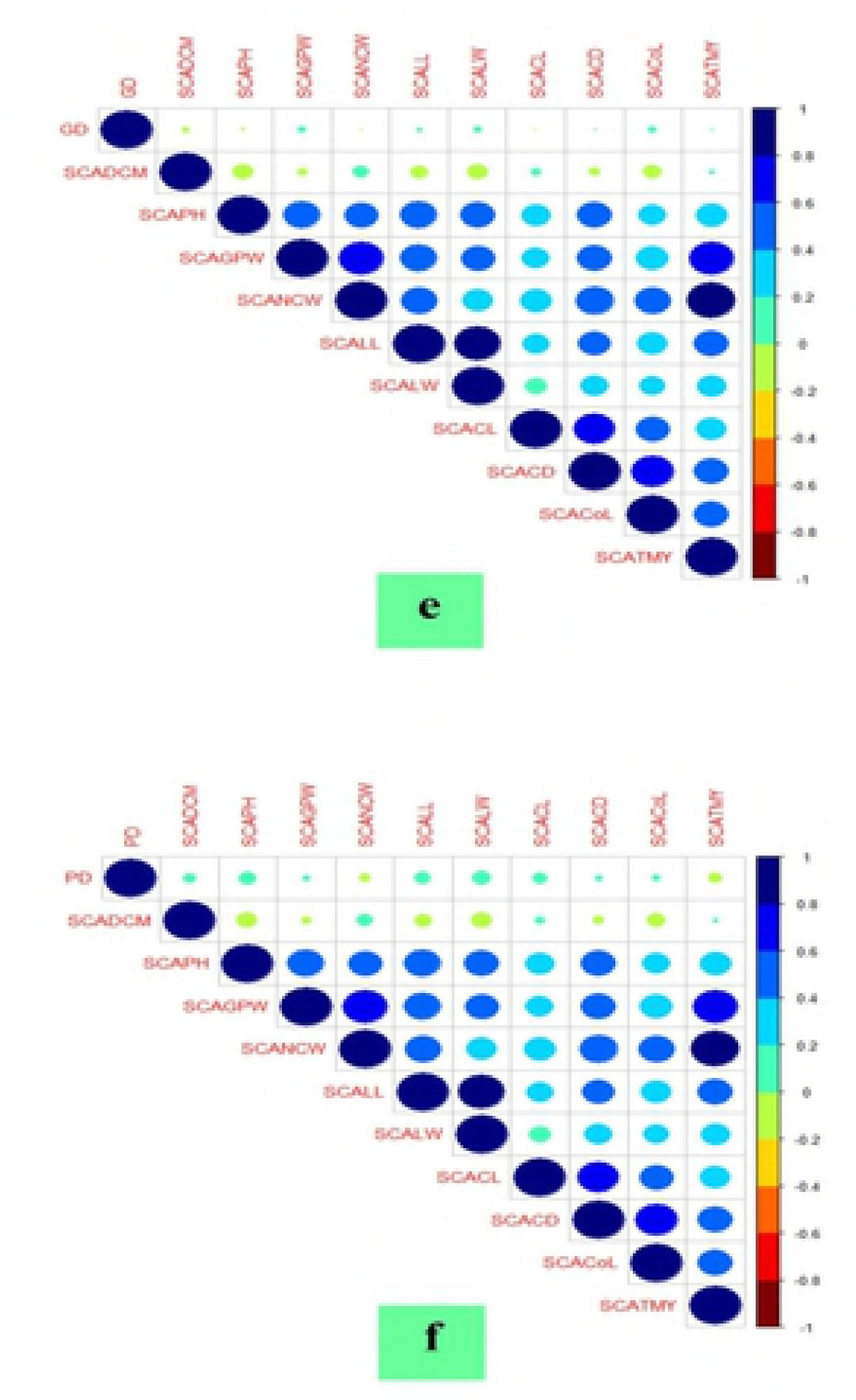
(e, f) e) PPMCC of GD with SCA, f) PD with SCA

**Fig 5.**
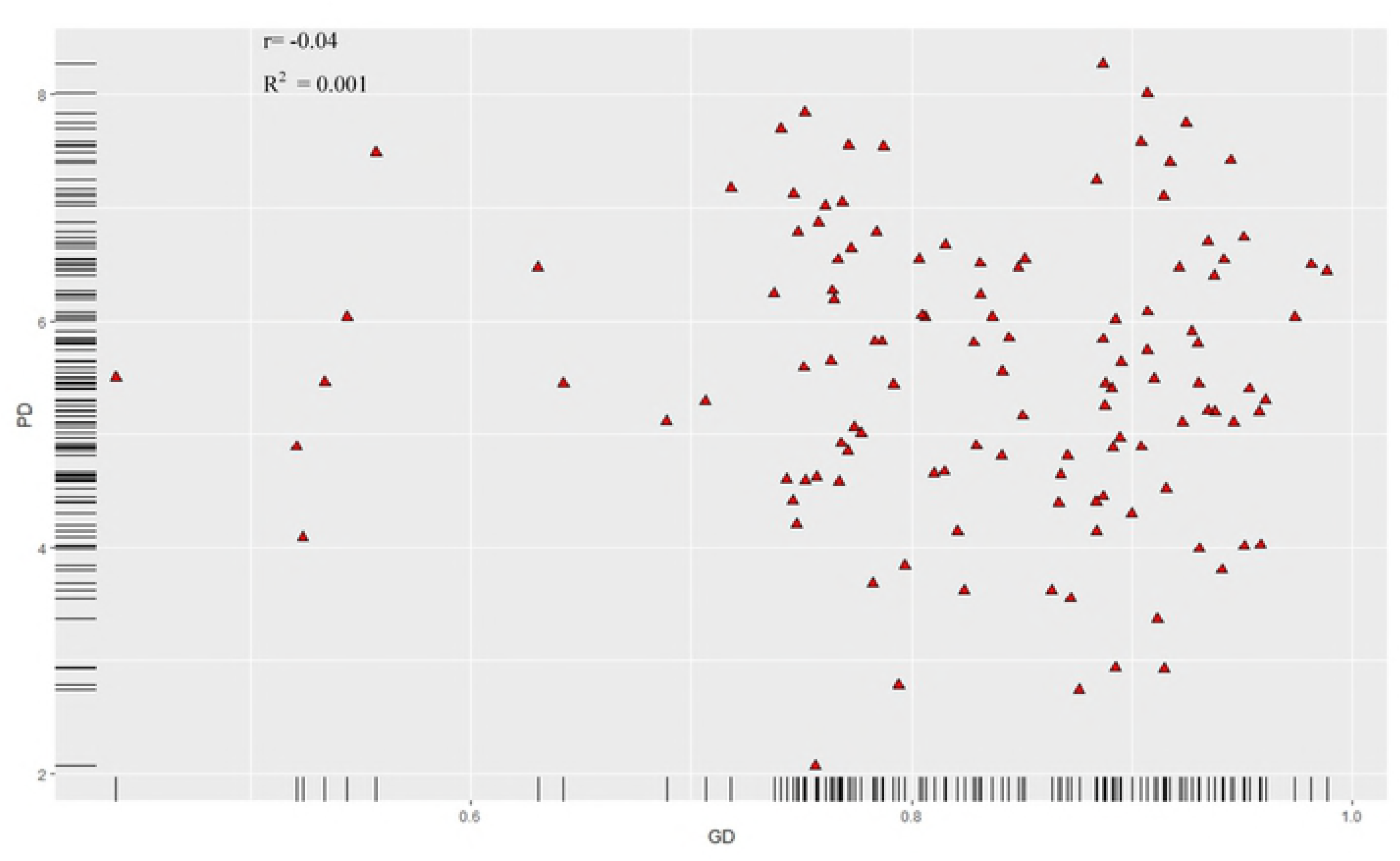
Relationship of phenotypic distance (PD) and microsatellite SSR, EST-SSRs based molecular distance (GD) based on all pair wise combinations of parental CMS lines and DH testers. GD is on X-axis and PD on Y-axis.

## Discussion

The biological phenomenon of heterosis has long been proved instrumental in enhancing agricultural productivity and has continuously captivated the plant breeders and geneticists to search on this unending process. The reliable prediction and identification of cross combinations of genetically diverse inbred lines giving heterotic performance is the key to successful hybrid breeding programmes, assuming positive correlation of GD and MPH [23]. The traditional approaches of quantitative genetics like, diallel analysis, generation mean analysis, line × tester analysis and estimating genetic components revealing various gene effects, are effective in unraveling genetic basis of heterosis [3, 41, 54, 69]. The line × tester analysis has revealed relative eligibility of lines and testers in determining heterotic testcross progenies. Then, GCA effects of parents, SCA effects of crosses along with estimation of genetic variance components and nature of gene action has enabled the selection of desirable parents and promising cross combinations to promulgate hybrid breeding across the crops including *Brassica oleracea* [13, 15, 69, 81]. The molecular markers based approaches for genetic analysis in prediction of heterosis include two ways viz. determining association between genetic divergence of parental lines and heterosis, and identification, mapping of heterotic QTLs displaying chromosome segments depicting heterosis [43]. Further, the allogamous breeding system and high inbreeding depression in *Brassica oleracea* vegetables, has facilitated the generation of numerous completely homozygous inbred lines through DH technology employing isolated microspore culture (IMC) [7-9, 25]. Globally, the genetic mechanism of CMS has been proved cost effective in enhancing the proportion of *Brassica* vegetables in hybrid vegetable seed industry. Thus, the global recognition of heterosis as ‘miraculous’ tool for feeding the world, paramount importance of CMS system in *Brassica* hybrid breeding and efficacy of DH technology in plant genetic studies, instigated us to determine heterotic crosses for agro-morphological, yield and commercial traits in cauliflower involving elite advance generation CMS lines and DH testers.

### Heritability, genetic components of variance, combining ability

The analysis of variance depicted highly significant differences among all the treatments for all the 16 agronomic traits, indicating considerable genetic differences among parents and their testcross progenies. And success of any crop breeding programme relies on genetic variation contained in studied germplasm. Similar results were reported by Garg and Lal [29], Verma and Kalia [81] for yield and related traits in early maturity Indian cauliflower, and for antioxidant traits in snowball cauliflower [69]. All the studied traits were found to be under the genetic control of both additive and non-additive gene effects, as revealed by significant mean squares of lines, testers and line × tester interactions (Table 3). The results are in agreement with Singh et al. [69] for antioxidant traits in cauliflower and Verma and Kalia [81] for days to 50% CM, leaf area, PH, MCW, NCW, curd compactness, GPW and HI in cauliflower using SI inbred lines. The analysis of genetic components of variance (Table 4) indicated the importance of SCA in developing heterotic crosses as revealed by higher value of σ^2^_sca_ than σ^2^_gca_ of lines and testers for majority of traits. Then, except days to 50% CM, all the vegetative and commercial traits showed predominance of dominance variance (σ^2^D) and greater than unity value of degree of dominance suggested over-dominance in the action of genes for vegetative and commercial traits in cauliflower. Further, except for days to 50% CM, the σ^2^_gca_/σ^2^_sca_ and σ^2^A/D suggested the non-additive genetic control of all the vegetative and commercial traits, and thus supported with high level of σ^2^_sca_, these results indicated the scope of heterosis breeding in genetic improvement of cauliflower with respect to these commercial traits. These findings are in accordance with Garg and Lal [29], Verma and Kalia [81] for curd and yield traits in cauliflower. As the response to natural and artificial selection relies on additive genetic variance, the narrow sense heritability (h^2^_ns_) holds a great promise in plant breeding as it provides basis to estimate accurate selection of genotypes based on phenotypic variance ascribed to additive genetic components [22]. In this study the low to intermediate level of h^2^_ns_ was observed for majority of vegetative and commercial traits suggesting non-additive genetic control of these traits, which might be due to large epistatic effects. We had also observed moderate estimates of h^2^_ns_ for antioxidant traits in cauliflower in previous study [69], then results are also in agreement with Xie et al. [85] for mineral content in Chinese cabbage. Thus, the early generation selection for these vegetative and commercial traits would be difficult due to dominance effects in the expression of phenotypic variance, and hence selection must be practiced in later generations. However, high h^2^_ns_ was observed with respect to earliness trait, days to 50% CM, and suggesting response to selection in early generation could be efficient. Further studies may be carried out in multiple standard environments for reaffirmation of these effects.

The combining ability analysis have been successfully utilized in crop breeding for evaluating parental performance and understanding dynamics of genes involved in trait expression. The parental GCA estimates in desirable direction also indicates potentiality of parents in generating promising breeding populations. In the present investigation, the significantly high GCA effects of parental lines in desirable direction for the respective vegetative and commercial traits are due to predominance of additive genetic effects of genes and additive × additive interactions [15, 69]. It depicts a desirable gene flow from parents to progeny at high frequency and these parental lines exhibiting high significant GCA for the respective traits in desirable direction can be utilized to stack favorable alleles via recombination and selection [1, 13, 15, 28, 69]. Further, our results revealed that none of the parents was good general combiner for all the studied vegetative and commercial traits. These findings are in conformity with results obtained by SI and CMS lines in cauliflower for yield and quality traits [13, 69, 81] and it suggested the requirement of multiple breeding programmes in suitable experimental and mating designs for the development of productive cultivars with the accumulation of positive alleles of genes. On the other hand, the parental lines depicting GCA in opposite direction for the respective traits can be utilized to generate desirable mapping population to study the genetics of respective traits [28]. The SCA, which reflects the loci having non-additive and epistatic gene effects, can be utilized to determine specific heterotic crosses for respective trait of interest. The significantly high SCA effects manifested in desirable direction by low × low testcrosses (poor GCA effects of both male and female parents) for instance Ogu307-33A × DH-18-8-3 for days to 50% CI, Ogu1A × DH-18-8-3 for days to 50% CM, Ogu22-1A × DH-53-6 for GPW and Ogu22-1A × DH-53-6 for NCW may be attributed to dominance × dominance type of interaction having especially complementary epistatic effects [24, 69]. This inconsistent association of GCA and SCA of respective crosses for respective traits is the indication of complex interaction of genes for quantitative traits [73]. Our results are corroborated by the findings of Verma and Kalia [81] for growth and yield traits in cauliflower using SI inbred lines and Singh et al. [69] for antioxidant traits using CMS lines. The majority of testcross progenies manifesting significantly high SCA in desirable direction had at least one of the parents reflecting poor GCA effects (poor × good general combiner or good × poor general combiner). The examples of such crosses are Ogu22-1A × DH-53-10 for days to 50% CM, Ogu122-1A × DH-18-8-3 for PH, Ogu12A × DH-53-10 for CoL, Ogu307-33A × DH-18-8-3 for GPW and it may be attributed to good combiner parent depicting favourable additive effects and poor combiner parent displaying epistatic effects [24, 69]. The crosses, manifesting significant SCA in desirable direction for respective traits, having both parents with good GCA (good general combiner × good general combiner) such as Ogu33A × DH53-1 for MCW, Ogu33A × DH-53-1 for NCW and Ogu33A × DH-53-1 for TMY, suggested the role of cumulative effects of additive × additive interaction of positive alleles [24, 69]. These findings are in compliance with results of Verma and Kalia [81] for growth and curd traits in cauliflower and Singh et al. [69] for antioxidant pigments in cauliflower. Concurrently, some of the crosses had poor SCA effects for the respective traits, despite involving parents with significant GCA, and it might be ascribed to absence of any interaction among the positive alleles of genes. Similar results were also reported by Singh et al. [69] with respect to quality traits in cauliflower and indicated the value of SCA in contrast to GCA in determining specific crosses superior for respective vegetative or commercial traits. Thus, our results suggested that breeders must pay attention to both GCA and SCA in the selection of elite parents for the development of heterotic hybrids. Further, the recombination breeding and random mating in conjunction with selection among segregates (recurrent selection), synthetics, composites, may be exploited to harness utility of both additive and non-additive gene effects in cauliflower [81]. The high SCA effects is not always correlated with significantly high heterosis and concurrently, the heterotic crosses exhibiting high MPH and BPH were not always had significant SCA effects. In the present study regarding this context, the heterotic crosses such as Ogu2A×DH-53-1, Ogu2A×DH-53-9 for days to 50% CI, Ogu33A×DH-53-9 for days to 50% CM, Ogu115-33A×DH-53-1 for GPW, Ogu118-6A×DH-53-1 for LSI, Ogu118-6A×DH-53-1 with respect to LW (Table 7) and crosses Ogu1A×DH-53-10 for CoL, Ogu126-1A×DH-53-1 for CL, Ogu115-33A×DH-53-6 for CD, Ogu33A×DH-53-1 for CSI, Ogu309-2A×DH-53-10 for MCW, Ogu309-2A×DH-53-10 with respect to NCW, Ogu126-1A×DH-53-10 for HI, Ogu309-2A×DH-53-10 for TMY (Table 8) were among top 10 crosses out of overall 120 crosses having significant MPH and BPH in desirable direction for the respective traits, but all these testcrosses had non significant poor SCA effects (Table 7, 8). Similar types of crosses were also reported for antioxidant traits in cauliflower [69]. It might be in response to the fact that the GCA of a parental line and SCA effects of a specific cross is dependent upon the particular lines, germplasm used in analysis, whereas heterosis is determined in response to mid parent, better parent or standard check.

### Cluster analysis, allelic diversity, and genetic structure

The study of morphological and molecular diversity is most vital in selecting desirable parents for hybrid breeding. The identification of heterotic pools and analyzing existing genetic variation in CMS lines is the preliminary requisite for efficient use of elite CMS lines in heterosis breeding. Study of genetic diversity at morphological and molecular level has been regarded as potential tool in identification of promising parental lines for developing heterotic hybrids in *Brassica oleracea* [17, 19, 58, 87]. Based upon PCA and HCA of 26 parental CMS lines and DH testers for 16 phenotypic traits, it was evident that all the parental lines had sufficient genetic variation. Then 55.1% variation was depicted by first two axes PC1 and PC2, thus PCA was efficient in determining genetic differentiation among parental lines. The PCA and NJ clustering based on molecular data represent better informative results for correct analysis and to be useful in crop improvement programme [17]. The PCA analysis and NJ clustering based on 87 SSR, EST-SSRs loci reaffirm that all the DH testers remained in two different sub-clusters of single group including two CMS lines Ogu125-8A and Ogu33-1A which showed close affinity with DH testers. Then CMS line Ogu125-8A with significantly high GCA in desirable direction for days to 50% CM, GPW, MCW, NCW, HI, TMY could be useful in developing high yielding early hybrid. The CMS line Ogu33-1A having significant GCA in desirable negative direction for earliness traits, days to 50% CI and days to 50% CM, could be used for generation of short duration early hybrids. Thus, the information pertaining to morphological and genetic diversity along with GCA could be useful in selecting desirable CMS lines as female parent for the development of cultivars with desirable traits. Similar types of findings have been reported by Dey et al. [17] and Parkash et al. [58] with respect to CMS lines in *Brassica oleracea*.

Further, we observed high allele frequency of overall 511 alleles through 87 genomic-SSR and EST-SSRs loci in 26 parental CMS and DH lines with average allelic frequency of 5.87 alleles per locus. It is quite high as compared to results reported by Parkash et al. [58], who observed only 58 total alleles with an average of 2 alleles per locus by 29 polymorphic SSRs in CMS lines of *Brassica oleracea,* and El-Esawi et al. [19], who reported 47 alleles with an average of 3.92 alleles per locus by 12 SSRs in *Brassica oleracea* genotypes. The quite high number of total alleles and high allelic frequency per locus revealed in the present investigation is quite possible as we used over all more numbers of genomic-SSR and EST-SSRs (total 350 microsatellite primers) distributed throughout the *Brassica oleracea* genome (n = 9, CC, 2n = 2x = 18), of which 87 loci depicted clear cut polymorphism. Of 350 microsatellites, > 50% was EST-SSRs primers, and EST-SSRs are derived from transcribed regions of genome and are having highly conserved sequences among homologous genes. They depicts the allelic diversity within or adjacent to genes and that might be more informative functionally and have higher transferability rate to related taxa in contrast to genomic SSRs [75, 80]. Then, we obtained high value of mean expected heterozygosity (*H_e_*), which is 0.68 indicating high genetic diversity in the studied genotypes as *H_e_* corresponds to genetic diversity. The PIC in genetic studies is utilized as a measure of informativeness of a marker locus for linkage analysis [19, 56] and it categorizes informative markers as highly informative (PIC ≥ 0.5), reasonably informative (0.5 < PIC >0.25) and slightly informative (PIC < 0.25) [19, 56]. In the present study the PIC content of 87 polymorphic loci ranged from 0.24-0.80 (Table 6), which classified all the 87 loci (g-SSR and EST-SSRs) as slightly informative (1 primer cnu107), reasonably informative (12 primers) and highly informative markers (74 primers) as per PIC content (Table 6), suggesting their ability in genetic differentiation of CMS and DH lines of cauliflower under study. The mean PIC content of 0.63 in present investigation based on 87 g-SSR and EST-SSRs was higher than the mean PIC of 0.316 observed for 165 cauliflower inbred lines by Zhu et al. [92] and 0.60 as recorded for 57 genotypes of *Brassica oleracea* comprising 51 cultivars of cauliflower by Zhao et al. [93]. The higher PIC value for most of loci revealed wide genetic diversity in the studied parental CMS and DH lines. The Bayesian genetic structure analysis based on posterior probability of data for a given k revealed 4 main sub-clusters of 26 parental CMS and DH lines at k = 4 with minor admixture. Cluster III mainly included all the DH testers along with two CMS lines Ogu125-8A and Ogu33-1A. Thus the 20 CMS lines were grouped into four clusters and maximum number of CMS lines was found in cluster I. These results suggested that DH testers were quite different genetically as compared to CMS lines. In all the clusters minor admixture was observed from each of clusters among themselves, which indicated the somewhat gene flow among the parental lines of different groups.

### Association of genetic distances and combining ability with heterosis

Numerous studies in different crops have been done to utilize the genetic distances in prediction of heterotic crosses (27, 34, 38, 40, 44, 73], assuming positive correlation of genetic distances with heterosis [23], but the correation of GD and heterosis is not absolute and significantly high level of heterosis can be obtained involving parents with low, intermediate or high genetic distance between them. Genetic distances based on both phenotypic and genotypic data are utilized to study the genetic variation among different genotypes or parental inbred lines. In the present investigation, high level of Euclidean distance (PD: 2.07 to 8.27) based on 16 phenotypic characters and GD (0.44-0.83) based on 87 genomic-SSR and EST-SSRs loci was reported among the CMS lines and DH testers of heterotic crosses. This might be due to the fact that CMS lines and DH testers used as female and male parent of testcross progenies were genetically quite dissimilar as reported by phenotypic and SSR, EST-SSRs based cluster analysis. The conflicting reports are available regarding correlations of genetic distances, heterosis and combining ability. In the present study, no correlation was observed between two distance measurements, based on morphological data (PD) and molecular data (GD). This is in contrary to the findings of Gupta et al. [33] who reported significantly positive correlation of GD and PD (r = 0.2) at P < 0.001 in pearl millets. Our results of no correlation between two distance measures might be due the fact that morphological traits showing continuous variation are largely influenced by environment and polygenic inheritance, linkage disequilibrium could result such relationship between two distance matrixes [10, 11, 33, 83]. Both the distance measures displayed no significant correlation with SCA of all the traits, suggesting genetic distances might not be effective in predicting SCA effects. Our results are in conformity with Su et al. [73] who also reported no significant association between genetic distances and SCA in chrysanthemum. However, Tian et al. [78], Lariepe et al. [44] reported significant correlation between total GD and SCA for length of terminal raceme in rapeseed, for grain yield and plant height in maize, respectively. Thus the association of GD with SCA is complicated. Further, our results suggested that SCA effects had stronger significant positive correlation with MPH and BPH for all the studied traits (0.52-0.83) at P ≤ 0.01. These results are in conformity with the findings of Zhang et al. [88], Su et al. [73], Tian et al. [78] in barley, chrysanthemum and rapeseed, respectively and indicated non-additive gene effects for heterosis. The GD and PD differed in their ability to predict MPH and BPH for different traits. Neither GD nor PD displayed any significant correlation with MPH and BPH for days to 50% CM, and CL. GD also exhibited no significant correlation with heterosis for LL and CoL. Similarly, PD showed no significant association with heterosis for majority of traits except LL. However, GD was significantly correlated with MPH and BPH for commercial traits viz. PH, GPW, NCW, LW, CD and TMY in desirable direction. These results are in line with the theory proposed by Falconer and Mackay [23]. In general, GD had greater magnitude of PPMCC than PD with heterosis for all the traits under study. The variability in correlation coefficients between hetrosis for respective traits and genetic distances may reflect allele numbers controlling the trait expression [34]. Wegary et al. [83] also highlighted the significant importance of GD in contrast to PD for predicting hybrid performance in maize. Our results are in agreement with the findings of Wegary et al. [83], who reported significant correlation of GD with heterosis for grain yield, plant height and ear height, similary of morphological distance with heterosis for certain traits in quality protein maize. The results obtained are also in line with the findings of Jagosz [36], who reported significant association of GD (based on RAP and AFLP markers) with heterosis for total and marketable yield in carrot. On the other hand, Tian et al. [78] and Su et al. [73] reported no significant correlation of PD and GD with MPH and BPH for any traits in rapeseed, chrysanthemum, respectively. Likewise, results are also in contrary with the findings of Geleta et al. [30] and Kawamura et al. [40] in pepper and chinese cabbage, respectively, suggesting no utility of GD in prediction of heterosis, while Krishnamurthy et al. [42] suggested selection of parents with intermediate divergence based on AFLP markers for getting more number of heterotic hybrids for yield in chilli using CMS lines. Regarding cole group of vegetables (*Brassica oleracea*), we only found a single report of describing interrelationships between genetic distances and heterosis, which is on broccoli (*B. oleracea* var. *italic* L.) by Hale et al. [34] using DH based population. They observed significantly negative correlation between total GD (based on SRAP, AFLP, SSR markers) and heterosis for all the traits, suggesting reduction in heterosis with the increase in genetic distances. Thus, our study is the first comprehensive report regarding interrelationships between GD (based on SSR, EST-SSRs) and heterosis for commercial traits in snowball cauliflower, suggesting significant correlation in desirable direction for respective traits. Hence, based on our results, we recommend the application of genomic-SSR and EST-SSRs based genetic distances in prediction of heterosis for yield and commercial traits involving CMS and DH based parental inbred lines in snowball cauliflower (*Brassica oleracea* var. *botrytis* L.). The non significant or poor correlation between GD and heterosis for certain traits might be due to lack of linkage between different alleles responsible for expression of particular trait and molecular marker used for estimating GD, inadequate coverage of entire genome, epistasis, DNA markers may be from unexpressed region of genome having no interaction with commercial traits and heterosis [5, 34, 53, 86]. The molecular marker based GD would be more predictive of heterosis, when there are strong dominance effects among hybrids, high heritability, linkage of molecular markers and QTLs of traits of interest [5, 34, 53, 86]. Hence, based on our results and previous findings by other researchers, it is quite evident that significance of genetic distances in prediction of heterosis inevitably depends upon, methods used to calculate genetic distances, type of molecular markers, genome coverage, region of genome, crop, breeding system, traits under consideration, type of germplasm and environmental conditions.

## Conclusions

In conclusion, our study is the first report on determining heterotic groups based on combining ability for morphological, yield and commercial traits using *Ogura* cybrid cytoplasm based CMS lines and DH testers. We also presented the first comprehensive report on predicting the association of genome wide EST-SSRs based GD and morphological traits based PD with heterosis, of F_1_ hybrids involving CMS and DH parental lines, for commercial traits in snowball cauliflower (*Brassica oleracea* var. *botrytis*). Analysis of variance of parents and their testcrosses revealed the presence of sufficient significant genetic variability, enabling the scope for crop improvement. Significant genetic differentiation was also observed among the parental CMS and DH lines using morphological and molecular markers. Present investigation also emphasizes the relevance of both GCA and SCA in the selection of elite parents for the improvement of yield and commercial traits and predicting appropriate breeding strategies for the crop genetic improvement, developing high yielding hybrids, synthetics and composites in cauliflower. Highly significant correlation of SCA with heterosis suggested the role of non-additive gene effects in heterosis. The findings of our study further suggested that genetic distances of SSR, EST-SSRs based molecular data can be used as reliable predictor of heterosis for commercial traits in CMS and DH based heterotic crosses of cauliflower. Although, the contrasting results obtained in different studies previously regarding efficacy of genetic distances in prediction of heterosis, invites further investigation with a different sets of large number of molecular markers covering entire genome, and different set of parental germplasm, in multiple standard environments.

## Acknowledgements

First author is thankful to Head ICAR-IARI, Regional Station, Katrain, (H.P.) for providing necessary facilities during research period and ICAR-IARI, New Delhi, during Ph.D. research programme. We also acknowledge the help of Dr. S. Dash, IASRI in statistical analysis using SAS software.

## Compliance with ethical standards

All the authors declare that they have no conflict of interest

## Author Contributions

**Conceptualization:** Saurabh Singh, S.S. Dey

**Data curation:** Saurabh Singh, S.S. Dey, Reeta Bhatia, Kanika Sharma

**Formal Analysis:** Saurabh Singh, S.S. Dey

**Investigation:** Saurabh Singh, S.S. Dey, Raj Kumar

**Resources:** S.S. Dey, Reeta Bhatia, Raj Kumar, T.K. Behera

**Software:** Saurabh Singh, S.S. Dey

**Supervision:** Raj Kumar

**Visualization:** S.S. Dey, T.K. Behera

**Writing original draft:** Saurabh Singh

**Review and editing:** S.S. Dey

**Supporting Information**

**S1 Table. List of 87 polymorphic genomic-SSR and EST-SSRs out of 350 microsatellite markers used for molecular diversity analysis.**

**S2 Table. Estimates of SCA effects of 120 test cross progenies for yield and horticultural traits.**

**S3 Table. Characterization of parental CMS and DH lines including commercial checks for 16 agronomic traits.**

**S4 Table. Estimates of phenotypic distance (PD), based on 16 phenotypic traits and genetic distance (GD), based on g-SSR, EST-SSRs molecular data, between parental lines and testers.**

## References

1. Arashida FM, Maluf WR, Carvalho RC. General and specific combining ability in tropical winter cauliflower. Hortic Bras. 2017; 35: 167–173. http://dx.doi.org/10.1590/s0102-053620170203.

2. Bayer PE, Golicz AA, Tirnaz S, Chan CK, Edwards D, Batley J. Variation in abundance of predicted resistance genes in the Brassica oleracea pangenome. Plant Biotechnol J. 2018; https://doi.org/10.1111/pbi.13015.

3. Bansal P, Banga S, Banga SS. Heterosis as investigated in terms of polyploidy and genetic diversity using designed Brassica juncea amphidiploids and its progenitor diploid species. PLOS ONE. 2012; 7: e29607.

4. Betran FJ, Ribaut JM, Beck D, de Leon DG. Genetic diversity, specific combining ability, and heterosis in tropical maize under stress and nonstress environments. Crop Sci. 2003; 43: 797–806.

5. Bernardo R. Relationship between single-cross performance and molecular marker heterozygosity. Theor Appl Genet. 1992; 83: 628–634.

6. Bhatia R, Dey SS, Sharma PC, Kumar R. In vitro maintenance of CMS lines of Indian cauliflower: an alternative for conventional CMS-based hybrid seed production. J Hortic Sci Biotechnol. 2015; 90: 171–179. https://doi.org/10.1080/14620316.2015.11513169.

7. Bhatia R, Dey SS, Sood S, Sharma K, Sharma VK, Parkash C, Kumar R. Optimizing protocol for efficient microspore embryogenesis and doubled haploid development in different maturity groups of cauliflower (Brassica oleracea var. botrytis L.) in India. Euphytica. 2016; 212: 439–454.

8. Bhatia R, Dey SS, Sood S, Sharma K, Parkash C, Kumar R. Efficient microspore embryogenesis in cauliflower (Brassica oleracea var. botrytis L.) for development of plants with different ploidy level and their use in breeding programme. Sci Hortic. 2017; 216: 83–92.

9. Bhatia R, Dey SS, Parkash C, Sharma K, Sood S, Kumar R. Modification of important factors for efficient microspore embryogenesis and doubled haploid production in field grown white cabbage (Brassica oleracea var. capitata L.) genotypes in India. Sci Hortic. 2018; 233: 178–187.

10. Burstin J, Charcosset A. Relationship between phenotypic and marker distances: theoretical and experimental investigations. J Hered. 1997; 79: 477–483.

11. Camussi A, Ottaviano E, Calinski T, Kaczmarek Z. Genetic distances based on quantitative traits. Genetics. 1985; 111: 945–962.

12. Cress CE. Heterosis of the hybrid related to gene frequency differences between two populations. Genetics. 1966; 53: 269.

13. Dey SS, Sharma SR, Bhatia R, Parkash C, Barwal RN. Superior CMS (Ogura) lines with better combining ability improved yield and maturity in cauliflower (Brassica oleracea var. botrytis). Euphytica. 2011; 182: 187–197.

14. Dey SS, Bhatia R, Sharma SR, Parkash C, Sureja AK. Effeccts of chloroplast substituted Ogura male sterile cytoplasm on the performance of cauliflower (Brassica oleracea var. botrytis L.) F_1_ hybrids. Sci Hortic. 2013; 157: 45–51.

15. Dey SS, Singh N, Bhatia R, Parkash C, Chandel C. Genetic combining ability and heterosis for important vitamins and antioxidant pigments in cauliflower (Brassica oleracea var. botrytis L.). Euphytica. 2014; 195: 169–181.

16. Dey SS, Bhatia R, Parkash C, Sharma S, Dabral M, Mishra V, Bhardwaj I, Sharma K, Sharma VK, Kumar R. Alteration in important quality traits and antioxidant activities in Brassica oleracea with Ogura cybrid cytoplasm. Plant Breed. 2017a; 136: 400–409.

17. Dey SS, Bhatia R, Bhardwaj I, Mishra V, Sharma K, Parkash C, Kumar S, Sharma VK, Kumar R. Molecular-agronomic characterization and genetic study reveals usefulness of refined Ogura cytoplasm based CMS lines in hybrid breeding of cauliflower (Brassica oleracea var. botrytis L.). Sci Hortic. 2017b; 224: 27–36.

18. Earl DA, vonHoldt BM. STRUCTURE HARVESTER: a website and program for visualizing STRUCTURE output and implementing the Evanno method. Conservation Genet Resour. 2012; 4: 359–361. https://doi.org/10.1007/s12686-011-9548-7.

19. El-Esawi MA, Germaine K, Bourke P, Malone R. Genetic diversity and population structure of Brassica oleracea germplasm in Ireland using SSR markers. C R Biol. 2016; 339: 133–140.

20. Esposito MA, Bermejo C, Gatti I, Guindon MF, Cravero V, Cointry EL. Prediction of heterotic crosses for yield in Pisum sativum L. Sci Hortic. 2014; 177: 53–62.

21. Evanno G, Regnaut S, Goude J. Detecting the number of clusters of individuals using the software STRUCTURE: a simulation study. Mol Ecol. 2005; 14: 2611–2620.

22. Evans LM, Tahmasbi R, Jones M, Vrieze SI, Abecasis GR, Das S, Bjelland DW, de Candia TR, Yang J, Goddard ME, Visscher PM, Keller MC. Narrow-sense heritability estimation of complex traits using identy-by-descent information. Heredity. 2018; https://doi.org/10.1038/s41437-018-0067-0.

23. Falconer DS, Mackay TFC. Introduction to quantitative genetics, 4th edn. Longman, London; 1996.

24. Fasahat P, Rajabi A, Rad MJ, Derera J. Principles and utilization of combining ability in plant breeding. Biom Biostat Int J. 2016; 4: 1–24. https://doi.org/10.15406/bbij.2016.04.00085.

25. Ferrie AMR, Mollers C. Haploids and doubled haploids in Brassica spp. for genetic and genomic research. Plant Cell Tiss Organ Cult. 2011; 104: 375–386.

26. Fujimoto R, Uezono K, Ishikura S, Osabe K, Peacock WJ, Dennis ES. Recent research on the mechanism of heterosis is important for crop and vegetable breeding systems. Breed Sci. 2018; 68: 145–158.

27. Fu D, Xiao M, Hayward A, Fu Y, Liu G, Jiang G, Zhang H. Utilization of crop heterosis: a review. Euphytica. 2014; 197: 161–173.

28. Gao W, Baars JJP, Dolstra O, Visser RGF, Sonnenberg ASM. Genetic variation and combining ability analysis of bruising sensitivity of Agaricus bisporus. PLOS ONE. 2013; 8: e76826. https://doi.org/10.1371/journal.pone.0076826.

29. Garg N, Lal T. Heterosis for growth and curd characters in Indian cauliflower (Brassica oleracea var. botrytis L.). Crop Improv. 2005; 32: 193–199.

30. Geleta LF, Labuschagne MT, Viljoen CD. Relationship between heterosis and genetic distance based on morphological traits and AFLP markers in pepper. Plant Breed. 2004; 123: 467–473.

31. Groszmann M, Greaves IK, Fujimoto R, Peacock WJ, Dennis ES. The role of epigenetics in hybrid vigour. Trends Genet. 2013; 29: 684–690.

32. Groszmann M, Gonzalez-Bayon R, Greaves IK, Wang L, Huen AK, Peacock WJ, Dennis ES. Intraspecific Arabidopsis hybrids show different patterns of heterosis despite the close relatedness of the parental genomes. Plant Physiol. 2014; 166(1): 265–80.

33. Gupta SK, Nepolean T, Shaikh CG, Rai K, Hash CT, Das RR, Rathore A. Phenotypic and molecular diversity-based prediction of heterosis in pearl millet (Pennisetum glaucum L. (R.) Br.). Crop J. 2018; 6: 271–281.

34. Hale AL, Farnham MW, Nzaramba MN, Kimbeng CA. Heterosis for horticultural traits in broccoli. Theor Appl Genet. 2007; 115: 351–360.

35. Hasan Y, Briggs W, Matschegewski C, Ordon F, Stützel H, Zetzsche H, Groen S, Uptmoor R. Quantitative trait loci controlling leaf appearance and curd initiation of cauliflower in relation to temperature. Theor Appl Genet. 2016; 129: 1273–1288.

36. Jagosz B. The relationship between heterosis and genetic distances based on RAPD and AFLP markers in carrot. Plant Breed. 2011; 130: 574–579.

37. Johnson HW, Robinson HF, Comstock RE. Estimation of genetic and environmental variability in soybeans. Agron J. 1955; 47: 314–318.

38. Kaushik P, Plazas M, Prohens J, Vilanova S, Gramazio P. Diallel genetic analysis for multiple traits in eggplant and assessment of genetic distances for predicting hybrids performance. PLOS ONE. 2018; 13: e0199943.

39. Kalinowski ST, Taper ML, Marshall TC. Revising how the computer program CERVUS accommodates genotyping error increases success in paternity assignment. Mol Ecol. 2007; 16: 1099–1106. http://dx.doi.org/10.1111/j.1365-294x.2007.03089.x.

40. Kawamura K, Kawanabe T, Shimizu M, Nagano AJ, Saeki N, Okazaki K, Kaji M, Dennis ES, Osabe K, Fujimoto R. Genetic distance of inbred lines of Chinese cabbage and its relationship to heterosis. Plant Gene. 2016; 5: 1–7.

41. Kempthorne O. An introduction to genetic statistics. John Wiley & Sons, Inc., New York; 1957.

42. Krishnamurthy SL, Rao AM, Reddy KM, Ramesh S, Hittalmani S, Rao MG. Limits of parental divergence for the occurrence of heterosis through morphological and AFLP marker in chilli (Capsicum annuum L.). Curr Sci. 2013; 104: 738–746.

43. Lariepe A, Mangin B, Jasson S, Combes V, Dumas F, Jamin P, Lariagon C, Jolivot D, Madur D, Fievet J, Gallais A, Dubreuil P, Charcosset A, Moreau L. The genetic basis of heterosis: multiparental quantitative trait loci mapping reveals contrasted levels of apparent overdominance among traits of agronomical interest in Maize (Zea mays L.). Genetics. 2012; 190: 795–811.

44. Lariepe A, Moreau L, Laborde J, Bauland C, Mezmouk S, Decousset L, Mary-Huard T, Fievet JB, Gallais A, Dubreuil P, Charcosset A. General and specific combining abilities in a maize (Zea mays L.) test-cross hybrid panel: relative importance of population structure and genetic divergence between parents. Theor Appl Genet. 2017; 130: 403- 417.

45. Lauss K, Wardenaar R, Oka R, van Hulten MHA, Guryev E, Keurentjes JJB, Stam M, Johannes F. Parental DNA methylation states are associated with heterosis in epigenetic hybrids. Plant Physiol. 2018; 176: 1627–1645.

46. Lin YR, Lee JY, Tseng MC, Lee CY, Shen CH, Wang CS, Liou CC, Shuang LS, Paterson AH, Hwu KK. Subtropical adaptation of a temperate plant (Brassica oleracea var. italica) utilizes non-vernalization-responsive QTLs. Sci Rep. 2018; 8: 13609. https://doi.org/10.1038/s41598-018-31987-1.

47. Li H, Chen X, Yang Y, Xu J, Gu J, Fu J, Qian X, Zhang S, Wu J, Liu K. Development and genetic mapping of microsatellite markers from whole genome shortgun sequences in Brassica oleracea. Mol Breed. 2011; 28: 585–596.

48. Li H, Yuan J, Wu M, Han Z, Li L, Jiang H, Jia Y, Han X, Liu M, Sun D, Chen C, Song W, Wang C. Transcriptome and DNA methylome reveal insights into yield heterosis in the curds of broccoli (Brassica oleracea L. var. italica). BMC Plant Biol. 2018; 18: 168. https://doi.org/10.1186/s12870-018-1384-4.

49. Lippman ZB, Zamir D. Heterosis: revisiting the magic. Trends Genet. 2007; 23: 60–66.

50. Liu X, Han F, Kong C, Fang Z, Yang L, Zhang Y, Zhuang M, Liu Y, Li Z, Lv H. Rapid introgression of fusarium wilt resistance gene into the elite cabbage line through the combined application of a microspore culture, genome background analysis, and disease resistance-specific marker assisted foreground selection. Front Plant Sci. 2017; 8: 354.

51. Lv H, Wang Q, Liu X, Han F, Fang Z, Yang L, Zhuang M, Liu Y, Li Z, Zhang Y. Whole-genome mapping reveals novel QTL cluster associated with main agronomic traits of cabbage (Brassica oleracea var. capitata L.). Front Plant Sci. 2016; 7: 989. https://doi.org/10.3389/fpls.2016.00989.

52. Maggioni L, von Bothmer R, Poulsen G, Lipman E. Domestication, diversity and use of Brassica oleracea L., based on ancient Greek and Latin texts. Genet Resour Crop Evol. 2018; 65: 137–159.

53. Makumbi D, Betran JF, Banziger M, Ribaut JM. Combining ability, heterosis and genetic diversity in tropical maize (Zea mays L.) under stress and non-stress conditions. Euphytica. 2011; 180: 143–162.

54. Melchinger AE, Piepho HP, Utz HF, Muminovic J, Wegenast T, Torjek O, Altmann A, Kusterer B. Genetic basis of heterosis for growth-related traits in Arabidopsis investigated by testcross progenies of near-isogenic lines reveals a significant role of epistasis. Genetics. 2007; 177: 1827–1837.

55. Meyer RC, Torjek O, Becher M, Altmann T. Heterosis of biomass production in Arabidopsis. Establishment during early development. Plant Physiol. 2004; 134: 1813- 1823.

56. Moges AD, Admassu B, Belew D, Yesuf M, Nijuguna J, Kyalo M, Ghimire SR. Development of microsatellite markers and analysis of genetic diversity and population structure of Colletotrichum gloeosporioides from Ethiopia. PLOS ONE. 2016; 11: e0151257. https://doi.org/10.1371/journal.pone.0151257.

57. Murray MG, Thompson WF. Rapid isolation of high molecular weight plant DNA. Nucleic Acid Res. 1980; 8: 4321.

58. Parkash C, Kumar S, Singh R, Kumar A, Kumar S, Dey SS, Bhatia R, Kumar R. ‘Ogura’- based ‘CMS’ lines with different nuclear backgrounds of cabbage revealed substantial diversity at morphological and molecular levels. 3Biotech. 2018; 8: 27. https://doi.org/10.1007/s13205-017-1047-4

59. Perrier X, Jacquemoud-Collet JP. DARwin software. 2006; http://darwin.cirad.fr/

60. Pritchard J, Stephens M, Donnelly P. Inference of population structure using multilocus genotype data. Genetics. 2000; 155: 945–959.

61. Robinson HS. Quantitative genetics in relation to breeding on the central of mendalism. Indian J Genet. 1966; 26: 171–187.

62. Rosen A, Hasan Y, Briggs W, Uptmoor R. Genome-based prediction of time to curd induction in cauliflower. Front Plant Sci. 2018; 9: 78.

63. RStudio Team. RStudio: Integrated Development for R. RStudio, Inc., Boston, MA, 2015; URL: http://www.rstudio.com/.

64. Samec D, Urlic B, Salopek-Sondi B. Kale (Brassica oleracea var. acephala) as a superfood: Review of the scientific evidence behind the statement. Crit Rev Food Sci Nutr. 2018; 20: 1–12. https://doi.org/0.1080/10408398.2018.

65. SAS Institute Inc. SAS Online Doc, Version 9.4. Cary, NC; 2013.

66. Sehgal N, Singh S. Progress on deciphering the molecular aspects of cell-to-cell communication in Brassica self-incompatibility response. 3Biotech. 2018; 8: 347. https://doi.org/10.1007/s13205-018-1372-2.

67. Sharma SR. Cabbage. In: Sirohi PS (ed) Vegetable crop production. Division of Vegetable Sciencce, IARI, New Delhi, 2003.

68. Singh S, Vidyasagar. Effect of common salt (NaCl) sprays to overcome the self-incompatibility in the S-allele lines of Brassica oleracea var. capitata L. SABRAO J Breed Genet. 2012; 44: 339–348.

69. Singh S, Bhatia R, Kumar R, Sharma K, Dash S, Dey SS. Cytoplasmic male sterile and doubled haploid lines with desirable combining ability enhances the concentration of important antioxidant attributes in Brassica oleracea. Euphytica. 2018a; 214: 207. https://doi.org/10.1007/s10681-018-2291-3.

70. Singh S, Dey SS, Bhatia R, Batley J, Kumar R. Molecular breeding for resistance to black rot [Xanthomonas campestris pv. campestris (Pammel) Dowson] in Brassicas: recent advances. Euphytica. 2018b; 214: 196. https://doi.org/10.1007/s10681-018-2275-3.

71. Sotelo T, Cartea ME, Velasco P, Soengas P. Identification of antioxidant-capacity related QTLs in Brassica oleracea. PLOS ONE. 2014; 9: e107290.

72. Su Y, Liu Y, Li Z, Fang Z, Yang L, Zhuang M, Zhang Y. QTL analysis of head splitting resistance in cabbage (Brassica oleracea L. var. capitata) using SSR and InDel markers based on whole genome re-sequencing. PLOS ONE. 2015; 10: e0138073. https://doi.org/10.1371/journal.pone.0138073.

73. Su J, Zhang F, Yang X, Feng Y, Yang X, Wu Y, Guan Z, Fang W, Chen F. Combining ability, heterosis, genetic distance and their intercorrelations for waterlogging tolerance traits in chrysanthemum. Euphytica. 2017; 213: 42.

74. Tams SH, Bauer E, Oettler G, Melchinger AE, Schon CC. Prospects for hybrid breeding in winter triticale: II. Relationship between parental genetic distance and specific combining ability. Plant Breed. 2006; 125: 331–336.

75. Taheri S, Abdullah TL, Yusop MR, Hanafi MM, Sahebi M, Azizi P, Shamshiri RR. Mining and development of novel SSR markers using next generation sequencing (NGS) data in plants. Molecules. 2018; 23: 399. https://doi.org/10.3390/molecules23020399.

76. Thakur AJ, Singh KH, Singh L, Nanjundan J, Khan YJ, Singh D. SSR marker variations in Brassica species provide insight into the origin and evolution of Brassica amphidiploids. Hereditas. 2018; 155: 6.

77. Thakur P, Vidyasagar, Singh S. Evaluation of cytoplasmic male sterile (CMS) progenies and maintainer lines for yield and horticultural traits in cabbage (Brassica oleracea var. capitata L.). SABRAO J Breed Genet. 2015; 47: 29–39.

78. Tian HY, Channa SA, Hu SW. Relationships between genetic distance, combining ability and heterosis in rapeseed (Brassica napus L.). Euphytica. 2017; 213: 1.

79. Teklewold A, Becker HC. Comparison of phenotypic and molecular distances to predict heterosis and F_1_ performance in Ethiopian mustard (Brassica carinata A. Braun.). 2006; 112 (4): 752–9. https://doi.org/10.1007/s00122-005-0180-3.

80. Varshney RK, Graner A, Sorrells MW. Genic microsatellite markers in plants: features and applications. Trends Biotechnol. 2005; 23: 48–55.

81. Verma VK, Kalia P. Combining ability analysis and its relationship with gene action and heterosis in early maturity cauliflower. Proc Natl Acad Sci, India, Sect B Biol Sci. 2017; 87: 877–884.

82. Wang W, Huang S, Liu Y, Fang Z, Yang L, Hua W, Yuan S, Liu S, Sun J, Zhuang M, Zhang Y, Zeng A. Construction and analysis of a high-density genetic linkage map in cabbage (Brassica oleracea L. var. capitata). BMC Genomics. 2012; 13: 523. https://doi.org/10.1186/1471-2164-13-523.

83. Wegary D, Vivek B, Labuschagne M. Association of parental genetic distance with heterosis and specific combining ability in quality protein maize. Euphytica. 2013; 191: 205–216.

84. Wei T, Simko VR. Package “corrplot”: Visualization of a Correlation Matrix (Version 0.84). 2017; https://github.com/taiyun/corrplot.

85. Xie F, Zha J, Tang H, Xu Y, Liu X, Wan Z. Combining ability and heterosis analysis for mineral elements by using cytoplasmic male-sterile systems in non-heading Chinese cabbage (Brassica rapa). Crop Pasture Sci. 2018; 69: 296–302.

86. Yu CY, Hu SW, Zhao HX, Guo AG, Sun GL. Genetic distances revealed by morphological characters, isozymes, proteins and RAPD markers and their relationships with hybrid performance in oilseed rape (Brassica napus L.). Theor Appl Genet. 2005; 110: 511–518.

87. Yousef EAA, Muller T, Borner A, Schmid KJ. Comparative analysis of genetic diversity and differentiation of cauliflower (Brassica oleracea var. botrytis) accessions from two ex situ gene banks. PLOS ONE. 2018; 13: e0192062. https://doi.org/10.1371/journal.pone.0192062.

88. Zhang X, Lv L, Lv C, Guo B, Xu R. Combining ability of different agronomic traits and yield components in hybrid barley. PLOS ONE. 2015; 10: e0126828. https://doi.org/10.1371/journal.pone.0126828.

89. Zhang X, Su Y, Liu Y, Fang Z, Yang L, Zhuang M, Zhang Y, Li Z, Lv H. Genetic analysis and QTL mapping of traits related to head shape in cabbage (Brassica oleracea var. capitata L.). Sci Hortic. 2016; 207: 82–88.

90. Zhao Z, Gu H, Sheng X, Yu H, Wang J, Huang L, Wang D. Genome-wide single-nucleotide polymorphisms discovery and high-density genetic map construction in cauliflower using specific-locus amplified fragment sequencing. Front Plant Sci. 2016; 7: 334.

91. Singh S, Singh R, Thakur P, Kumar R. Phytochemicals, functionality and breeding for enrichment of cole vegetables (Brassica oleracea L.). In: Petropoulos SA, Ferreira ICFR, Barros L, editors. Phytochemicals in vegetables: a valuable source of bioactive compounds. Bentham Science Publishers, UAE. 2018. Pp. 256–295.

92. Zhu S, Zhang X, Liu Q, Luo T, Tang Z, Zhou Y. The genetic diversity and relationships of cauliflower (Brassica oleracea var. botrytis) inbred lines assessed by using SSR markers. PLOS ONE. 2018; 13: e0208551. https://doi.org/10.1371/journal.pone.0208551

93. Zhao Z, Gu H, Sheng X, Yu H, Wang J, Zhao J, et al. Genetic diversity and relationships among loose-curd cauliflower and related varieties as revealed by microsatellite markers. Sci Hortic. 2014; 166: 105–110

